# A wave of WNT signalling balanced by secreted inhibitors controls primitive streak formation in micropattern colonies of human embryonic stem cells

**DOI:** 10.1101/440602

**Authors:** I. Martyn, A.H. Brivanlou, E.D. Siggia

## Abstract

Long range signalling by morphogens and their inhibitors define embryonic patterning yet quantitative data and models are rare, especially in humans. Here we use a human embryonic stem cell “gastruloid” system to model formation of the primitive streak (PS) by WNT. In the pluripotent state E-CADHERIN (E-CAD) transduces boundary forces to focus WNT signalling to colony border. Following application of WNT ligand, E-CAD mediates a wave of epithelial-to-mesenchymal (EMT) conversion analogous to PS extension in an embryo. By knocking out the secreted WNT inhibitors active in our system, we show that DKK1 alone controls the extent and duration of patterning. The NODAL inhibitor CERBERUS1 acts downstream of WNT to refine the endoderm versus mesoderm fate choice. Our EMT wave is a generic property of a bistable system with diffusion and a single quantitative model describes both the wave and our knockout data.

## Introduction

The WNT, BMP, and ACTIVIN/NODAL signalling pathways play a dominant and largely conserved role in vertebrate development. Despite extensive knowledge of the intracellular aspects of these pathways, however, many quantitative aspects of how they create and mediate large-scale patterns remain unknown, especially in the context of human development. A prime example of this is the patterning of the human epiblast, where in a period of approximately two days, and on a millimeter length scale, these three pathways overlap and create the necessary instructions to form and pattern the embryonic germ layers and future anterior-posterior axis of the human embryo. While mouse studies have shed some light on the involvement of these pathways in the patterning processes, differences in architecture, timing, and juxtaposition of extraembryonic and embryonic tissues makes the direct comparison difficult. Because of these differences and because the interactions of the aforementioned pathways are complex, with multiple morphogens and secreted inhibitors from overlapping regions acting at the same time, there is a need for an assay that allows for their control in a model human epiblast that also permits a dynamic readout on the scale of single cells.

We have previously shown that human embryonic stem cells (hESCs) confined to micropatterned colonies of 1 millimeter diameter can be used as *in vitro* assay to model the human epiblast^1,2^. These micropatterns self-organize in response to WNT, BMP, and ACTIVIN/NODAL signalling and recapitulate the patterning of germ layers observed during mammalian gastrulation. For example, stimulation with BMP4 for 48 hours results in concentric rings corresponding to ectoderm, mesoderm, endoderm, and extraembryonic tissue arranged from the center to edge. As current guidelines prohibit the studies of human embryos after 14 days (the 14 day rule) these models currently remain the best alternative to direct *in vivo* studies. More importantly, these models allow single-cell quantification and control over geometry, density, signalling strength, and genetics.

In subsequent work we exploited this assay to uncover how the BMP pathway contributes to this patterning^3^. Briefly, cells in the colonies are apically-basally polarized and BMP4 receptors are located on the baso-lateral sides of the cells, restricting access to the apically supplied medium except near the edges. This receptor mediated prepattern, already present in the pluripotent state, is reinforced by the secreted BMP inhibitor NOGGIN, which in humans is directly induced by BMP4. Due to diffusion and boundary conditions, NOGGIN is highest in the center and lowest at the colony edges, resulting in an effective gradient of BMP response across the colony.

We have also shown that BMP4 induces WNT ligand and that this WNT is necessary for the mesoderm and endoderm part of the pattern^4^. Additionally, we have shown that WNT signaling is sufficient to induce the self-organization of a human PS at the edge of colonies, and that co-presentation of WNT3A and ACTIVIN leads to the induction of functional human organizer cells that can induce an ectopic secondary axis in chick embryos. Since here again there is a self-organized patterned response to a uniformly presented ligand, our system offers an ideal environment to explore how WNT signalling leads to spatial organization, and specifically how the human PS forms and is spatially confined.

Here we address the molecular mechanisms underlying WNT-mediated self-organization of human PS. We show that two primary factors control patterning: E-CADHERIN (E-CAD) and DKK1. First, E-CAD establishes a pre-pattern by limiting the initial WNT response to the boundary. Secondly, and in parallel to the NOGGIN dynamics in the BMP case, the secreted inhibitor DKK1 is upregulated by a combination of WNT and NODAL signalling and is required to ultimately confine the PS to the colony boundary. Multiple single and double combinations of homozygous CRISPR/Cas9 knockouts of secreted inhibitors of the WNT and NODAL pathways were made and they confirmed that only DKK1 plays a major role in the spatial restriction of the PS. We found that CERBERUS1 (CER1) is also highly upregulated by a combination of WNT and NODAL signalling, but that in our cells it functions as a NODAL inhibitor rather than dual WNT/NODAL inhibitor. CER1 thus does not influence the size of the PS, but instead serves to bias the mesoderm versus endoderm fate decision in this region. We found also that in our DKK1^-/-^ cells E-CAD not only establishes a pre-pattern, but, via its mutual antagonism with WNT, generates a cooperative EMT wave that travels from the micropattern periphery to the center.

## Results

### WNT response is edge and density dependent and apically-basally symmetric

We previously showed that uniform application of WNT3A ligand to hESC micropatterns is sufficient to self-organize a PS-like structure, with mesoderm and endoderm emerging from an EMT on the colony periphery after 48 hours and with ACTIVIN/NODAL level biasing the choice of endoderm versus mesoderm^4^ (Figure 1A). We will use the union of SOX17 and BRA to define the spatial extent of the induced PS, and their ratio to define the proportion of endoderm versus mesoderm throughout. We also showed previously that despite the uniform application of WNT, the interior of the colony remains pluripotent, expressing both NANOG and SOX2^4^. This pattern, with pluripotent cells on the interior and mesoderm and endoderm cells on the periphery, represents the terminal spatial pattern we seek to understand.

**Figure 1.**
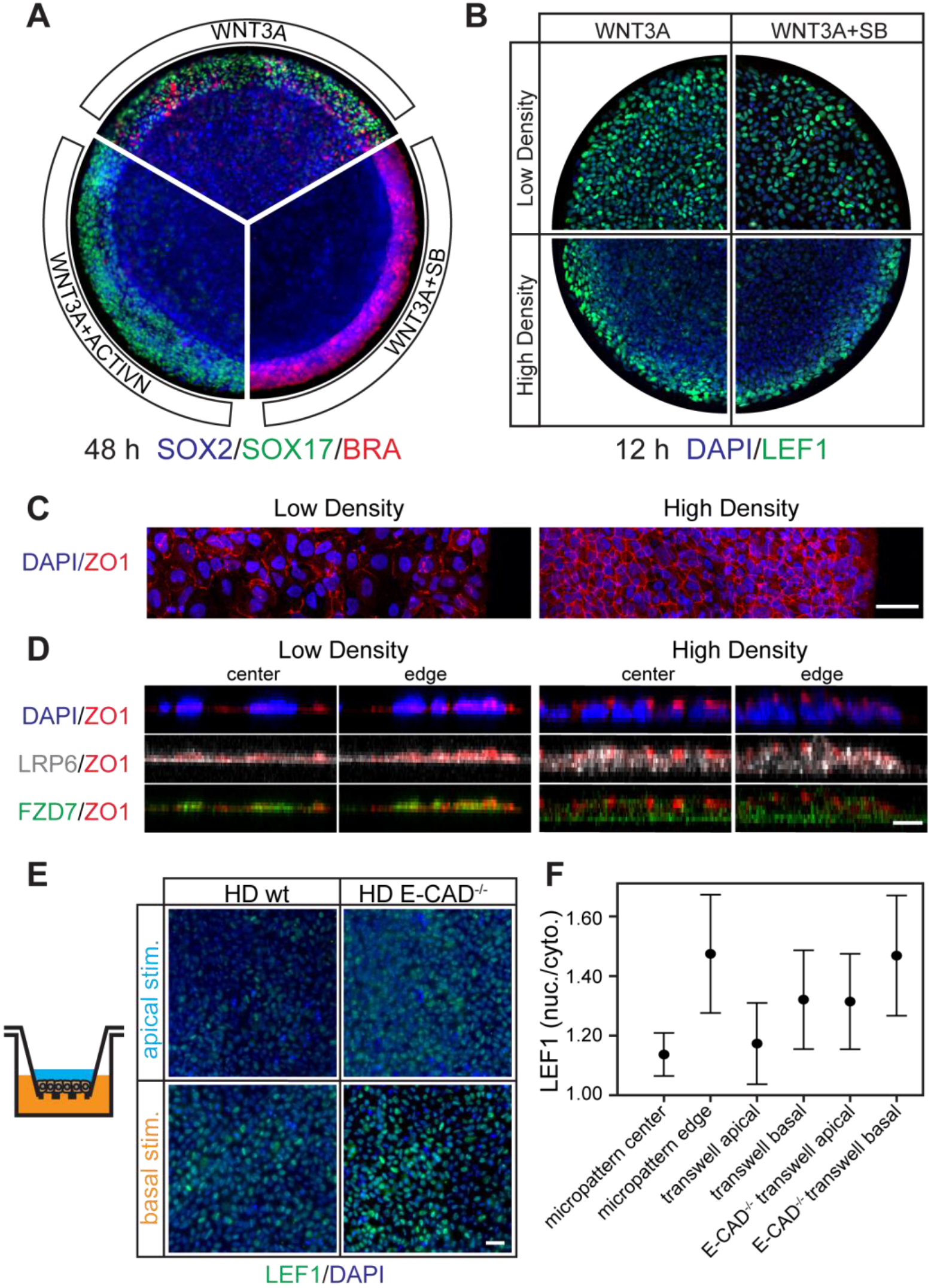
WNT3A response is edge and density dependent. (A) Pie sections of representative 1,000μm diameter micropatterned hESC colonies stimulated with WNT3A, WNT3A+SB, or WNT3A+ACTIVIN and fixed and stained after 48h for germ layer markers. All micropattern experiments were performed on at least three separate occasions with similar results, and unless mentioned otherwise, all other micropatterns shown are 1,000 μm in diameter (B) Micropatterns stimulated with WNT3A or WNT3A+SB at high density (22h after seeding, 3474 ± 430 cells/mm^2^) or low density (8h after seeding, 1810 ± 236 cells/mm^2^) and fixed and stained for LEF1 at 12h. (C) Maximum intensity projection of tight junction marker ZO1 and nuclear marker DAPI in low density and high density micropatterns immediately prior to stimulation. Note that the network of tight junctions is only fully formed in the high density micropatterns. Scale bar, 50μm. (D) Cross-sections showing the apical-basal position of WNT receptors relative to DAPI and ZO1 (apical marker). At high density the cells are in an apically-basally polarized epithelial state, as judged by the relative position ZO1 and DAPI. While not polarized themselves, the majority of the WNT receptors in this state do lie underneath the tight junctions, and so presumably are not be as accessible from the apical side as the basal side. This is in contrast to the low density state where cells are not epithelized and receptors are visible on both sides of ZO1. Additionally there is no significant edge-vs-center expression of the receptors at either density. Scale bar, 10μm. (E) and (F). Basally stimulated high density hESCs in transwell filters show a higher WNT3A response than when apically stimulated. However, judging by the quantification of LEF1 nuclear to cytoplasmic ratio in (F), this difference is not enough to explain the edge-vs-center difference in micropatterns. It takes the knockout of E-CAD and basal stimulation to reach the micropattern edge level of WNT activation. The error bars in (F) represent the standard deviation on 5,000 cells and the scale bar in (E) is 50μm.

In order to decipher the molecular mechanism underlying this spatial pattern, we first attempted to use the detection of nuclear β-CATENIN (β-CAT) as an early readout for canonical WNT signalling^5–7^. However, commercially available antibodies did not have adequate resolution on our dense epithelia (Figure S1A), so instead we used LEF1, a co-factor of β-CAT that is a direct target of WNT signalling with the same response profile as AXIN2 (Figure S1B) and is localized to the nucleus^8,9^. We found that the LEF1 response profile depended on colony density, with nuclear expression throughout the colonies at low density and restriction to the periphery at high density (Figure 1B). For the sake of comparison, the SMAD1 response due to BMP stimulation of micropatterns in this high density regime is also edge restricted. Co-presentation of the SMAD2 pathway inhibitor SB-431542, together with WNT3A did not change the outcome, demonstrating that the density dependence of the LEF1 pattern is specifically due to WNT, and not caused by secondary ACTIVIN/NODAL signaling.

As one of the factors involved in BMP4- induced self-organization was due to polarized signal reception, we first examined the localization of the WNT receptors^10^ FRIZZLED and LRP5/6 in our micropatterns. We find that while some WNT-receptors were detected on the apical side, they were predominantly and homogenously located basolaterally underneath the tight junctions (Figure 1D). We also find little distinction between edge and center. To functionally, test for signal reception, we cultured cells on transwell-filter culture dishes, where cells can be selectively stimulated from the apical or basal side. We cultured them at the same density as the high density micropatterns and stimulated them with WNT3A for 12 hours. While a stronger response was detected with basal rather than apical stimulation, the mean basal response on filters, however fell below the edge response on colonies (Figure 1E and F), suggesting that additional factors were involved in setting up the WNT response on the colony boundary.

### E-CAD knockdown sensitizes cells to WNT

Given the fact that E-CAD is classically considered to antagonize WNT signalling via its binding and sequestration of β-CAT^11–14^, and the fact that cells on the periphery of our micropatterns have fewer neighbouring cells and so presumably fewer E-CAD junctions, we hypothesized that E-CAD may contribute to the early WNT pattern. To test this hypothesis we made clonal CRISPR/Cas9 E-CAD knockout cell lines. We found that E-CAD^-/-^ hESCs could be passaged and seeded normally, grew at the same rate as wild type cells, maintained pluripotency markers, and still apical-basally polarized at high density to form intact epithelia (Figure S2A-D). E-CAD^-/-^ WNT receptor localization was also indistinguishable from wild type (Figure S2E).

Strikingly, however, the early WNT pattern was abolished in E-CAD^-/-^ micropatterns (Figure 2C). Cells at the center of high density colonies now showed nuclear localization of LEF1. Quantifying our results over multiple micropatterns, we observed that the WNT response is modestly biased to the edge, and generally comparable to the level in low density colonies (Figure 2D). Our transwell filter assay still showed an apical-basal asymmetry, but now the basal response was elevated to match the edge response of the parental cells (Figure 1E and F). These results demonstrate that the early WNT pattern is primarily due to E-CAD activity, and a minor influence exerted by WNT receptor accessibility.

**Figure 2.**
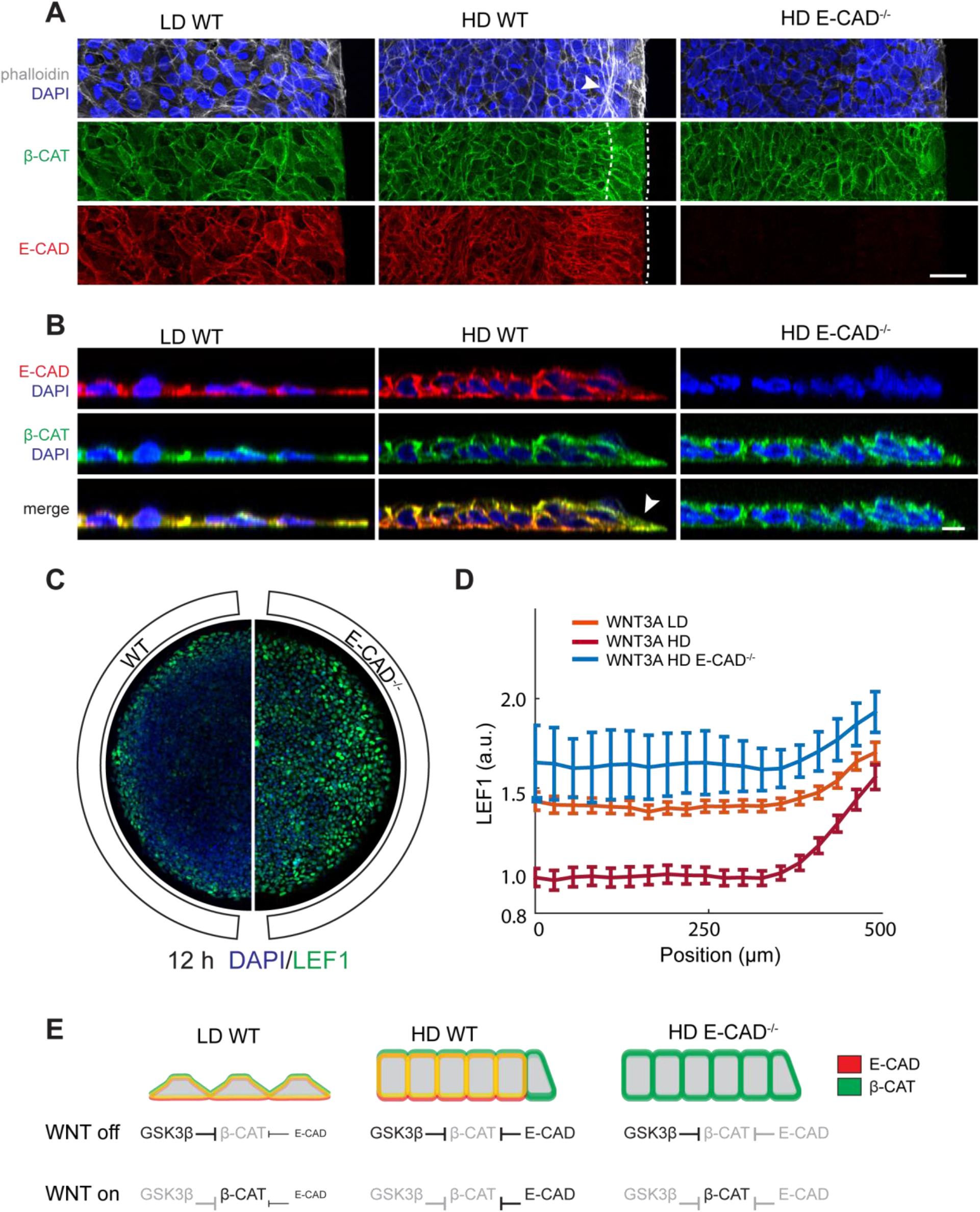
E-CAD knockdown eliminates early WNT3A pattern. (A) Maximum intensity projection of DAPI, actin (marked via phalloiden), β-CAT, and E-CAD in low density wild type, high density wild type, and high density E-CAD^-/-^ micropatterns immediately prior to WNT3A stimulation. Note the thick actin stress fibers (arrow) and cytoplasmic β-CAT (between the two dashed lines) only apparent on the edge of the high density wild type micropatterns, in the same region that shows the highest LEF1 response to WNT3A. Note also that outer facing side of cells on the micropattern edge are lower in E-CAD compared with sides of the same cells that that join with neighbouring cells (bottom dashed line). Scale bar, 50μm. (B) Cross-section of micropatterns from (A) showing the overlap of E-CAD and β-CAT. In low density wild type micropatterns there is no significant asymmetry in E-CAD or β-CAT localization (superposition is yellow), but at high density one can see unmatched free β-CAT (green) on cells on the periphery of the micropattern (arrow). Scale bar, 10μm. (C) 12h WNT3A response measure by LEF1 in high density wild type and E-CAD^-/-^ micropatterns. Knockdown of E-CAD allows WNT3A response to penetrate into the center of the micropattern. (D) Quantification (C) and comparison to the low density wild type micropatterns. Single cell expression data was binned radially and averaged. The final radial profile represents the average of *n*=25 colonies. Error bars here and on all following graphs represents the standard deviation among colonies. (E) Our results are consistent with a model where E-CAD acts as an additional negative inhibitor of β-CAT. At low density this inhibition is negligible and so WNT3A stimulation and down regulation of the β-CAT destruction complex results in a uniform upregulation of β-CAT and a LEF1 response. At high density, however, where there are more mature cadherin junctions, this inhibition is significant, and the pre-existing edge-vs-center asymmetry in E-CAD state result in a WNT induced pattern when the destruction complex is downregulated.

If all cells express E-CAD why are there spatial differences in the WNT response? We hypothesized that spatial differences in E-CAD localization or in the state of E-CAD junctions and their binding partners could account for spatial WNT signalling differences. E-CAD and β-CAT stains support this hypothesis as they show that in high density wild type micropatterns E-CAD is reduced in cells on the periphery and there is observable cytoplasmic β-CAT here as well (Figure 2A and B). Actin stress fibers have also been observed on hESC micropattern boundaries^15,16^ and have been implicated in E-CAD dysregulation^17^. Phalloiden staining in our wild type micropatterns revealed that there were indeed actin stress fibers and they were restricted to the region that is WNT responsive. Furthermore, these stress fibers are absent from low density or E-CAD^-/-^ micropatterns (Figure 2A), which suggests a connection between mechanics and WNT signalling.

### Disruption of E-CAD/β-CAT binding or actin cytoskeleton also sensitizes cells to WNT

To further test whether the classical connection between E-CAD, β-CAT, and the actin cytoskeleton was responsible for our early WNT pattern we performed two additional experiments. In the first we inserted into our E-CAD^-/-^ cells either constitutively expressed full length E-CAD or constitutively expressed E-CAD that lacked the β-CAT binding domain^18^ into the AAVS1 locus using TALENS. Clonal lines were cultured in micropatterns and stimulated for 12 hours to examine the WNT response. We found that the constitutively expressed full length E-CAD rescued the edge restricted phenotype but that the E-CAD without the β-CAT binding domain did not (Figure 3B). This shows that E-CAD binding to β-CAT is essential for the early WNT pattern.

**Figure 3.**
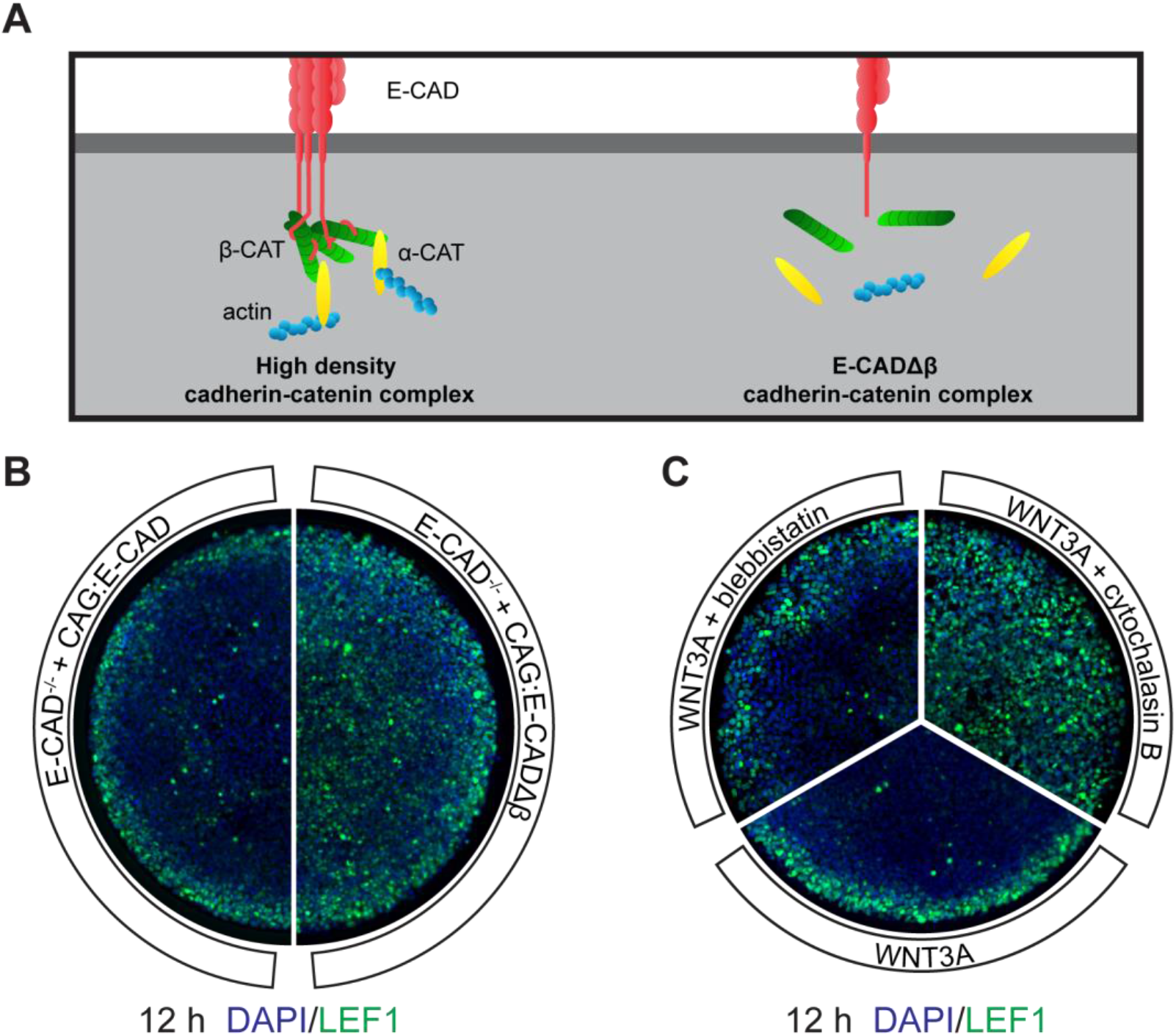
Other perturbations to E-CAD-β-CAT-Actin connections also eliminate the early WNT3A pattern. (A) Cartoon showing the conventional model of how E-CAD, β-CAT, and the actin cytoskeleton connect to one another. (B) E-CAD^-/-^ cells rescued with constitutively expressed full length E-CAD or E-CADΔβ were put in high density micropatterns and stimulated with WNT3A for 12h and then stained for LEF1. The full length E-CAD micropatterns recovered the wild type phenotype whereas those with E-CADΔβ did not, showing that the β-CAT link to E-CAD is essential for the 12h WNT pattern. (C) Wild type high density micropatterns stimulated with WNT3A, WNT3A+blebbistatin, or WNT3A+cytochalasin B, and stained for LEF1 after 12h. Blebbistatin blocks myosin II controlled actin contraction and cross-linking, whereas cytochalasin B more directly interferes with the actin cytoskeleton by reducing actin polymerization. Corresponding to this difference of degree of perturbation, we see a minor increase in the width of the LEF1 region with blebbistatin, and a more dramatic elimination of the LEF1 edge restriction with cytochalasin B.

In the second experiment we disrupted the actin cytoskeleton across the entire micropattern with the small molecule inhibitors blebbistatin or cytochalasin B while stimulating with WNT3A. Blebbistatin has been shown to dislodge E-CAD from the membrane into the cytoplasm in hESCs in pluripotency^19^, and cytochalasin B has an even more direct action on dissociating the actin cytoskeleton. We found that both reduced the edge restriction, with blebbistatin broadening the size of the LEF1 band and cytochalasin B permitting a WNT response even in the center of the colony (Figure 3C). Taken together, our results demonstrate that colony geometry acts via the cytoskeleton and E-CAD to bias WNT signalling to the colony boundary.

### Self-organization of the PS to the edge occurs in E-CAD^-/-^ cells

Having understood the reasons for the edge asymmetry in the initial response to WNT, we were surprised to see that the location of the PS was virtually the same in wild type and E-CAD^-/-^ colonies at 48 hours (Figure 4A). Examination of LEF1, SOX2 and BRA expression at intermediate times showed that a homogenous early expression of these markers gradually becomes restricted to the edge (Figure 4B). These dynamics are suggestive of a WNT induced secreted inhibitor that is highest in the center and progressively restricts WNT activity to the boundary.

**Figure 4.**
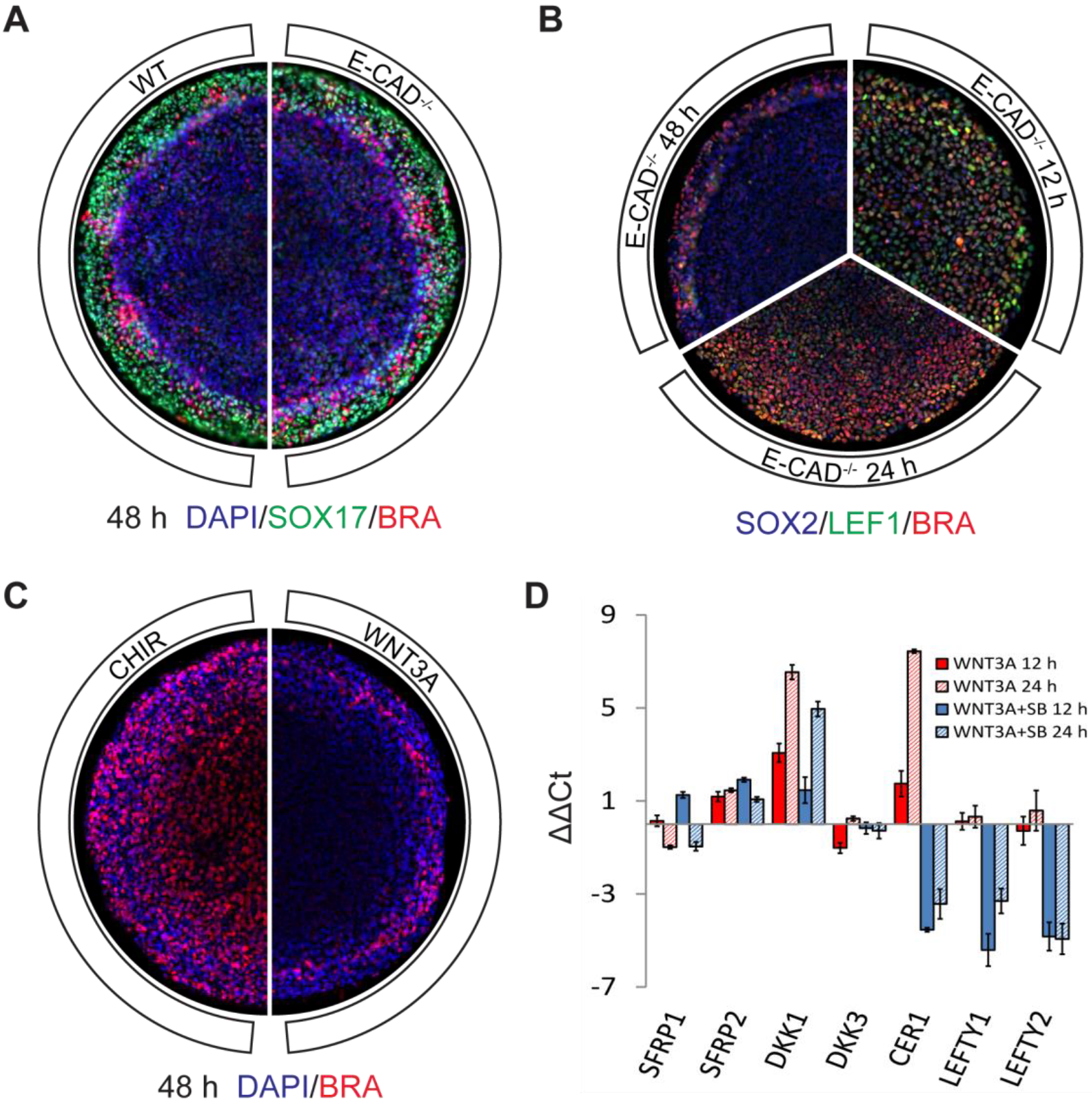
E-CAD not enough to explain long term WNT patterning, WNT3A also induces WNT and NODAL inhibitors. (A) High density wild type and E-CAD^-/-^ micropatterns fixed and stained for indicated markers after 48h of WNT3A stimulation. (B) Time-course of LEF1 and BRA expression in high density E-CAD^-/-^ micropatterns. Note that as time progresses the signalling response is gradually excluded from the colony center. (C) High density E-CAD^-/-^ micropatterned colonies stimulated with 6μM CHIR or WNT3A and fixed and stained for BRA after 48h. The dramatic difference between the two indicates that extra-cellular regulation of the WNT3A pathway may be a dominant factor, since CHIR is a small molecule and acts intercellularly, skipping extracellular regulation. (D) qPCR of secreted inhibitors of interest upon 12 and 24h of WNT3A or WNT3A+SB stimulation in micropatterned colonies. Note that the NODAL inhibitors are all severely downregulated when one inhibits the NODAL receptor with SB. (Note also that Conditioned Media also has endogenous ACTIVIN/NODAL activity that would contribute to the effect seen with SB even though no additional ACTIVIN was added).

We had previously shown that BMP4 directly induced the expression of its own inhibitor NOGGIN, which in turn was necessary and sufficient to restrict BMP signalling to the colony edge after 48 hours of stimulation^3^. To address if a similar mechanism of WNT3A inducing its own inhibitor was involved in this case as well, we activated the pathway with CHIR-99021, a small molecule compound that acts cell autonomously and will skip receptors and secreted inhibitors. After 48 hours stimulation the compound edge restriction was abolished (Figure 4C). This result strongly suggests the involvement of secreted inhibitors in WNT-mediated self-organization and PS formation.

### WNT induces WNT and NODAL inhibitors

To further test this hypothesis, and identify the relevant inhibitors, we focused on WNT inhibitors whose loss-of-function leads to early gastrulation defects phenotypes in the mouse. These include SFRP1, SFRP2, DKK1, and DKK3^20^. Additionally, as WNT activates NODAL^4^, and these the two ligands have been shown to act synergistically to induce mesendodermal genes^21^, we also included LEFTY1, LEFTY2, and CER1 on our list. qPCR was used to assess the induction of these inhibitors when cells were treated with WNT3A alone, or WNT3A+SB to distinguish direct versus indirect induction. After 12 and 24 hours of stimulation, expression of SFRP1, SFRP2, and DKK3 remain relatively unchanged regardless of WNT3A or WNT3A+SB treatment (Figure 4E). DKK1 expression, however, was highly up-regulated in response to WNT3A. Similar to what has been previously reported^21^, this appears to depend on synergy between the WNT and NODAL pathways, since DKK1 induction is lower in WNT3A+SB conditions. A stronger dependence on SMAD2 signaling was observed for the WNT3A induction of CER1 expression. Finally, the expression of LEFTY1 and LEFTY2 depend even more on NODAL signalling since they are also down-regulated in WNT3A+SB and cannot be activated with WNT3A alone. Thus DKK1 and CER1 emerged as the leading candidates to be involved in WNT self-organization.

### DKK1 controls the size of the PS

To test whether DKK1 and CER1 are required for the late time WNT pattern, we used CRISPR/Cas9 to generate DKK1 and CER1 knockouts. Each clonal line was stimulated with WNT3A and compared to wild type. After 12 hours of stimulation, DKK1^-/-^ colonies showed no difference with the control (Figure 5A). After 48 hours of treatment, however, the size of the PS was dramatically increased when compared to the wild type and E-CAD^-/-^ lines, with only a small center of SOX2+ undifferentiated cells remaining (Figure 5B and C). This result, which was confirmed in two additional clonal DKK1^-/-^ lines (Figure S4A), demonstrates that DKK1 activity is required for WNT-mediated patterning, and is consistent with a reaction-diffusion model.

**Figure 5.**
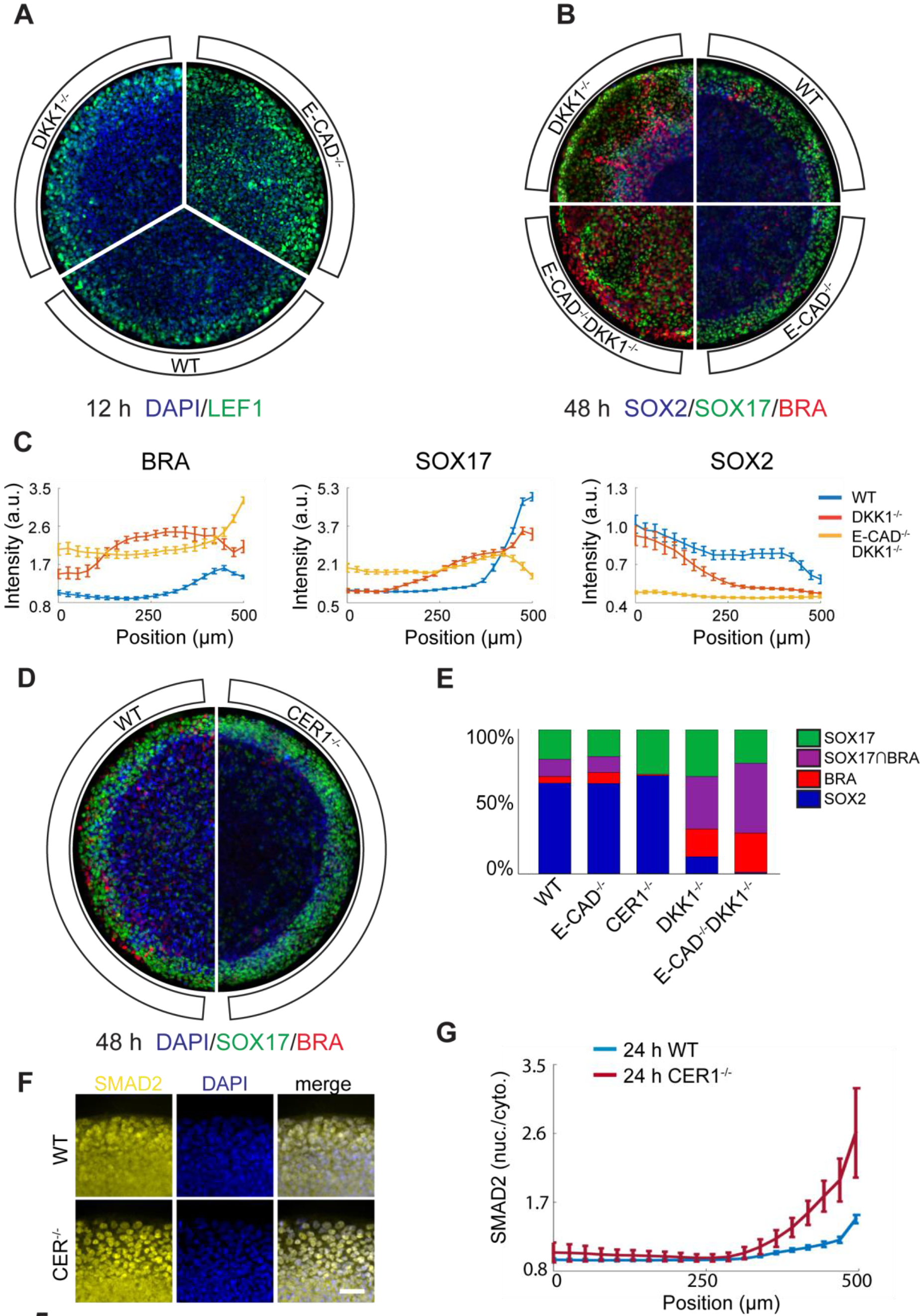
DKK1 controls spatial extent of WNT3A patterning and CER1 influences the mesoderm vs endoderm fate decision. (A) Comparison of 12h WNT3A response in high density micropatterns of DKK1^-/-^ cells vs E-CAD ^-/-^ cells vs wild type cells. As expected (since no DKK1 is transcribed in pluripotency) DKK1^-/-^ micropatterns resemble wild type micropatterns and are less sensitive to WNT3A than E-CAD^-/-^ cells. (B) After 48h of WNT3A stimulation on high density micropatterns, DKK1^-/-^ cells show a dramatic increase in WNT3A patterned region compared with E-CAD^-/-^ and wild type cells. DKK1 and E-CAD also act synergistically, as loss of both genes cause complete loss in the edge restriction of WNT3A patterning. (C) Quantification of IF for pluripotency marker SOX2 and differentiation markers SOX17 and BRA in DKK1^-/-^, E-CAD^-/-^, and DKK1^-/-^E-CAD^-/-^ cells (n=20 colonies per condition) following B. (D) Comparison of wild type and CER1^-/-^ micropatterns after 48h of WNT3A stimulation. Notice the higher number of BRA cells in the wild type. (E) Classification of cells from wild type, CER1^-/-^, DKK1^-/-^, E-CAD^-/-^, and DKK1^-/-^E-CAD^-/-^ micropatterns into 4 different subpopulations. Classification was performed by fitting single cell SOX17, BRA, and SOX2 levels to a Gaussian mixture model (See Methods and Figure S4G). Note that in addition to increasing the spatial extent of WNT3A patterning, DKK1 also influences the proportion of differentiated cells that commit to either mesoderm (BRA) or endoderm (SOX17), with significantly more cells expressing BRA when DKK1 is knocked out. Note also the decline in BRA cells in the CER1^-/-^ micropatterns compared to the wild type. (F) SMAD2 levels and the ring of activity are increased in CER1^-/-^ cells compared to wild type (micropatterns stimulated with WNT3A, fixed and stained after 48h). Scale bar, 50μm. (G) Quantification of F, *n*=20 colonies per condition.

To test for epistasis between WNT inhibition at early times mediated by E-CAD, and at late times by DKK1, we generated a double E-CAD^-/-^DKK1^-/-^ knockout line. In response to WNT3A stimulation for 48 hours, all cells in the micropatterned colonies differentiated, with no SOX2+ cells left in the center (Figure 5B and C). This suggests that DKK1 and E-CAD are the two major players among the collection of WNT inhibitors that block differentiation in our gastruloids.

Comparison of the expanded PS in the single and double knockouts with wild type also established that both the total number and the ratio of BRA to SOX17 cells changed (Figure 5E). Whereas the proportion BRA+/SOX17- cells in RUES2 wild type line was ~10%, in DKK1^-/-^ cells it doubled to 20%, and in E-CAD^-/-^DKK1^-/-^ it tripled to ^~^30%. The fraction of cells that express both BRA and SOX17 also greatly increases in the mutant lines. This suggests that in addition to determining the size of the PS, DKK1 may also be involved in the segregation of mesodermal and endodermal fates.

### CER1 biases mesoderm versus endoderm fate decision

When stimulated with WNT3A, CER1^-/-^ cells did not show any change in the size of the PS domain in comparison to wild type (Figure 5D). There was, however, a significant shift in the proportion of mesodermal versus endodermal fates. Unlike the DKK1^-/-^ or E-CAD^-/-^DKK1^-/-^ cells, this time the shift was towards greater endoderm, with almost all differentiated cells expressing SOX17 and none expressing BRA (Figure 5E). This represents a similar phenotype to that shown for WNT3A+ACTIVIN treatment (Figure 1A), and prompted us to investigate the status of NODAL/ACTIVIN signaling. We find that SMAD2 signalling is enhanced in the CER1^-/-^ knockout line (Figure 5F and G). It is known in mouse that CER1 inhibits BMP and NODAL signalling but not WNT signalling^22^ (at odds with other vertebrates systems where its tri-functionality motivates its name^20^). For human however it is only known that CER1 inhibits NODAL and a subset of BMP ligands, with a verdict on WNT inhibition still awaiting^23,24^. Since the size of the PS remains unchanged while SMAD2 signaling increases along with the proportion of endodermal cells in the CER1^-/-^ colonies, our results suggest that in hESCs CER1 acts primarily as a NODAL inhibitor rather than a WNT inhibitor.

As it was previously shown in the mouse that the most dramatic CER1 phenotype is when it is doubly knocked out with LEFTY1^25^, we also generated CER1^-/-^LEFTY1^-/-^ and LEFTY1^-/-^ clonal cell lines. We detected no difference in the WNT response phenotype between wildtype and LEFTY1^-/-^ cells, and no difference between CER1^-/-^ and CER1^-/-^LEFTY1^-/-^ cells (Figure S4B and C). In order to check for all other players identified in our RNA-seq and qPCR results, we generated DKK3^-/-^ and SFRP1^-/-^SFRP2^-/-^ clonal cell lines. None of these lines displayed any phenotypic difference when compared to wildtype (Figure S4D-F). We conclude that DKK1 and CER1 are the major secreted inhibitors that control WNT patterning in our gastruloids.

### An edge to center WNT/EMT wave in human gastruloids

The size of the PS region in the DKK1^-/-^ cell line at 48 hours is intermediate between the smaller wild type PS region and the fully converted PS region of the double E-CAD^-/-^DKK1^-/-^ cell line. Given that in both RUES2 and DKK1^-/-^ colonies WNT signalling begins at the edge (Figure 5A), an important and relevant question is whether the 48 hour result is at steady-state, or instead if given more time the PS would eventually expand inward and consume the entire colony. To address this question we fixed and stained wild type and DKK1^-/-^ micropatterns at 12, 24, 48, and 72 hours. We find that while differentiation starts similarly for both, the wild type micropatterns seems to reach a steady state by
24-48 hours while differentiation and EMT in the DKK1^-/-^ colonies continues to proceeds inwards, eventually almost consuming the entire micropattern by 72 hours (Figure 6A). This is consistent with a wave of WNT differentiation proceeding from the outer edge to the center.

**Figure 6.**
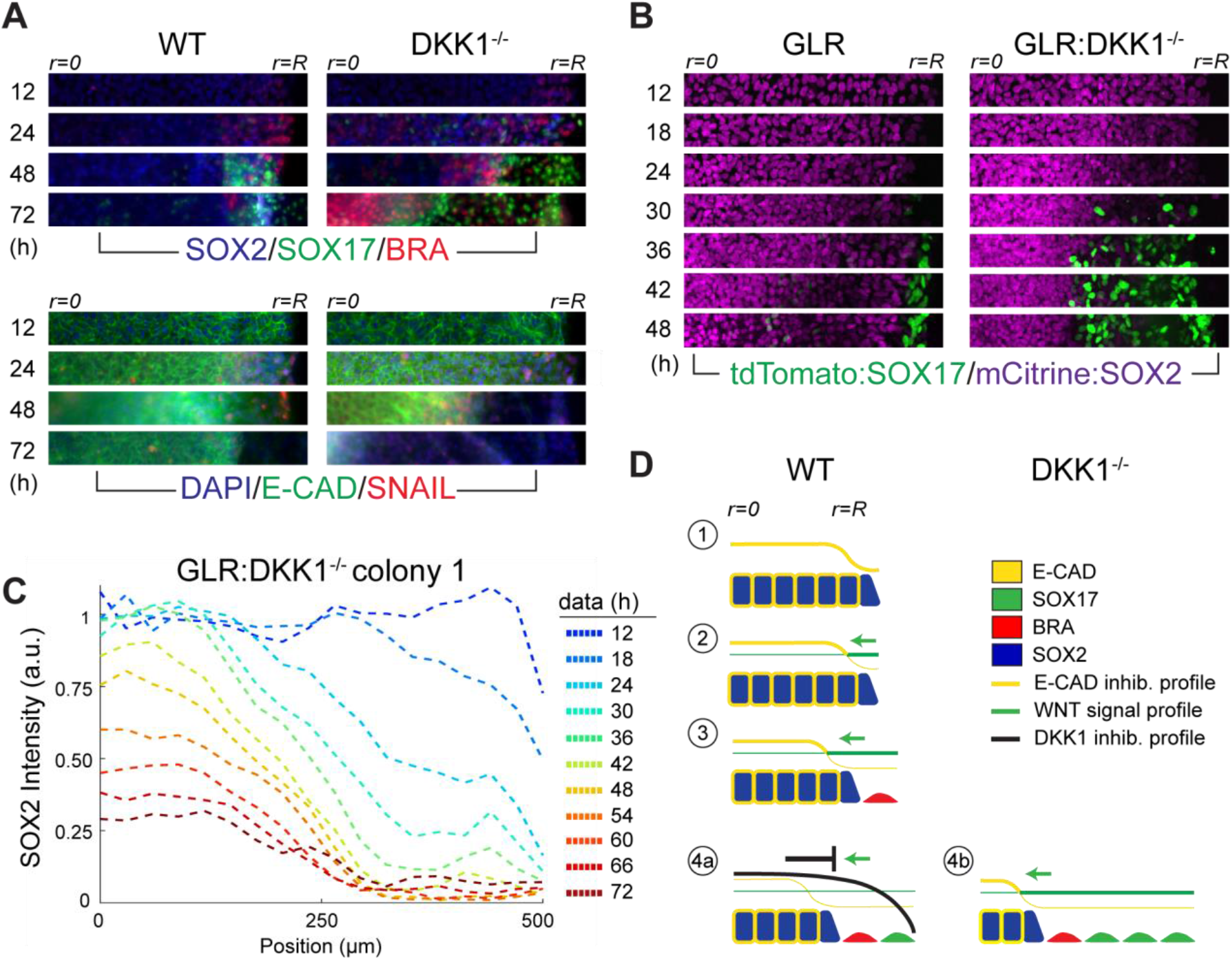
Patterning via a WNT/EMT wave. (A) Radial sections of WNT3A stimulated high density wild type or E-CAD^-/-^ micropatterns fixed and stained for the indicated markers at indicated times. Interior of each colony (r=0) is on the left of each section, the edge of the colony (r=R) is on the right. (B) Time-lapse radial sections of WNT3A stimulated high density GLR micropattern and GLR:DKK1^-/-^ micropattern. Ten micropatterns for each condition were imaged in the same session, and the examples shown here are representative. One notes that SOX17 turns on slightly earlier in the GLR:DKK1^-/-^ micropatterns than in the GLR micropatterns, and that, as with the immunostaining data, a wave of SOX2 downregulation and differentiation starts on the periphery. This wave halts in the GLR line, but continues proceeding inward in the GLR:DKK1^-/-^ micropatterns. (C) Quantification of single-cell SOX2 expression in the same GLR:DKK1^-/-^ micropattern shown in (B). (D) Qualitative model of WNT/EMT wave spreading and stabilization. ➀: prior to WNT3A stimulation E-CAD creates a bias so that only cells on the immediate periphery are sensitive to WNT ligand. ➁: Application of WNT3A ligand results in only boundary cells responding and differentiating. ➂: As these cells undergo EMT, they lose E-CAD junctions and expose interior cells, enabling them now to respond to the WNT ligand. ➃: If checked by secreted DKK1 from the differentiating cells, however, the boundary cells become protected from WNT ligand and the wave stops, as illustrated in (a); if left unchecked, this cycle will enable a wave of differentiation to travel progressively across the colony from outside to inside, as illustrated in (b).

To confirm the existence of this wave in our gastruloids and study it further, we knocked out DKK1 in a previously described RUES2-GLR (Germ Layer Reporter) cell line where ectoderm (SOX2), mesoderm (BRA), and endoderm (SOX17) germ layers are tagged with 3 separate fluorescent markers^4^. This enabled us to evaluate the change in differentiation and fate acquisition in the same micropattern across different times. Stimulation of the control RUES2-GLR with WNT3A leads to a downregulation of SOX2 and an upregulation of SOX17 that begins at the edge but stops a few cell layers inward, maintaining the PS at the periphery. RUES2-GLR colonies in which the DKK1^-/-^ mutation is introduced, however, display a wave of progressive downregulation of SOX2 and upregulation of SOX17, which, as in the stained time-course, begins at the outer edge and does not stop (Figure 6B and C).

What is the mechanism for such a wave? Given that E-CAD expression recedes as the differentiation front advances (Figure 6A) we posit that the wave results from a positive EMT feedback loop. WNT3A is known to induce EMT and to down-regulate E-CAD through the transcription factor SNAIL^26,27^. As illustrated in Figure 6D, WNT3A first down-regulates E-CAD in WNT susceptible edge cells and causes them to go through EMT. In so doing, these cells destabilize their E-CAD junctions with their neighbours. This leads to a domino like propagation of EMT from cell to cell via shared cell contacts. If the differentiated cells were induced to secrete a diffusing and thus long-ranged WNT inhibitor, i.e. DKK1, then it will accumulate in the center more than the edges and the wave would be expected to halt^3^. This is how the wild type cells achieve the steady state we observe, and why the DKK1^-/-^and GLR:DKK1^-/-^ cells fail to do so.

In addition to these dynamics, previous results have shown that WNT ligand also upregulates WNT production in hESCs^4^. To test if this endogenous WNT signalling contributed to the dynamics we compared wild type and DKK1^-/-^ micropatterns with and without IWP2, a small molecule that blocks all WNT ligand secretion. We find no significant differences (Figure S5A), most likely due to the fact that we were already stimulating our micropatterns with a high dose if WNT in the media and thus are in a saturated regime where endogenous WNT does not make any significant contribution to the dynamics. We also tested the involvement of ACTIVIN/NODAL signalling (which has a baseline activity in our media) in this wave by comparing DKK1^-/-^ micropatterns stimulated either with WNT3A or WNT3A+SB. Consistent with other studies of EMT^28^, our wave stops when we block ACTIVIN/NODAL signalling with SB (Figure S5B).

### A quantitative dynamic model

The spatial pattern of WNT signalling in our colonies is defined by inhibition from E-CAD at the earliest times and DKK1 at later times. These inhibitors operate very differently in space: E-CAD bridges adjacent cells, while DKK1 diffuses across the colony and also leaks out at the edges^3^. Downstream of WNT, NODAL and CER1 are produced and together with WNT generate mesoderm and endoderm fates. To fully unravel the complexity involved in this process we formulated a quantitative dynamic model.

A good model will use a portion of the data to fit parameters and then make testable informative predictions about the remainder of the data, and do this with as few variables as possible. With these criteria in mind we define a model where the intracellular WNT signal is normalized to [0,1] and where a simple Michaelis-Menten system of equations links it to the two inhibitors (see Supplemental Methods). One advantage of our formulation is that it is separable, so we can fit the DKK1 specific parameters to the E-CAD^-/-^ data and vice versa (Figure 7A). Since we cannot directly measure WNT levels we use the immunofluorescence data for LEF1 at 12 hours and the percentage loss of SOX2 at 48 hours as surrogates (Supplemental Methods).

**Figure 7.**
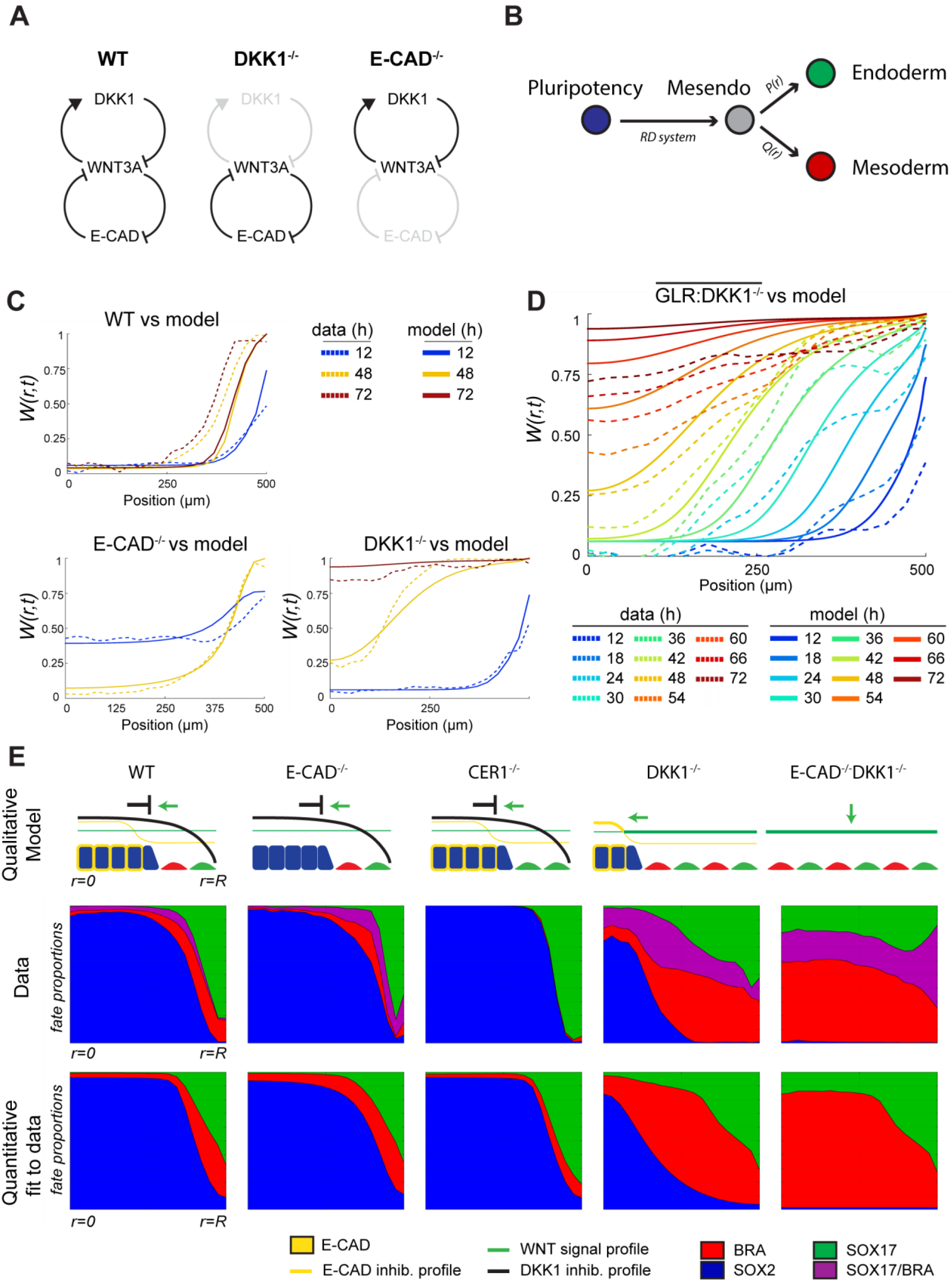
A quantitative model of WNT patterning dynamics in the PS. (A) Gene regulatory network of WNT3A, E-CAD, and DKK1 used in the model. Note how the DKK1^-/-^ and E-CAD^-/-^ cell lines can be used to simplify this network and fit the subcomponents separately. (B) Simple model of mesoderm versus endoderm fate decision. The reaction-diffusion (RD) system determines the probability that a cell at a given radius in a given background commits to differentiate by going from the pluripotent state to the intermediate mesdendo state. Once there, the probability *P*(*r*), which is a linearly rescaled function of radial nuclear SMAD2 profile, is used to determine the probability of cell going to endoderm versus mesodederm. (C) Comparison of the RD model to the WT, DKK1^-/-^, and E-CAD^-/-^ data. The fit was made using the 12h LEF1 response and the 48h differentiated cell response (i.e. BRA/SOX17 or 1-SOX2) in the DKK1^-/-^ and E-CAD^-/-^ micropatterns in Figure 5B after linearly rescaling the profiles to the interval [0,1]. The remaining data are predictions. (D) WNT/EMT wave in the DKK1^-/-^ micropatterns. Solid lines are the model predictions, dashed lines are the average of 10 continuously imaged GLR:DKK1^-/-^ micropatterns. (E) Qualitative diagram of WNT patterning dynamics for each genetic background, and fit of quantitative model to the data. For the data, single cells from the 48h micropatterns of each genetic background were classified into 4 different subpopulations according to a Gaussian mixture model based on SOX17, BRA, and SOX2 levels (see Methods). Classified cells were then further subdivided according to their radial position and the population was normalized, allowing us thus to obtain the probability of belonging to a specific fate at a specific radii given a specific genetic background.

Figure 7C shows the quality of the fits to the data at 12 and 48 hours in the two knockout lines that we consider quite acceptable (Suppl. Figures M2-3). Our model with no additional adjustments is then able to predict and reproduce the 72 hour data as well as the EMT wave, which is most visible in the GLR:DKK1^-/-^ cell line (Figure 7D). This live data is somewhat variable due to density variation and phototoxicity (see Supplemental Methods) but the prediction of the shape of the wave front and where it should be at a specific time closely follows the observations. The collapse of the wave after it reaches a radius of about 200 μm is also predicted and the model agrees with both the live data and the immunofluorescence data in Figure 6A.

The model makes explicit and mathematically precise that the reciprocal inhibition between WNT and E-CAD (Figure 7A) can give rise to bistability, and how, in the absence of the long-range secreted inhibitor DKK1, the bistability resolves by an inward propagating wave that eliminates the epithelial state in favour of the PS mesenchyme. The model situates the WNT system within a general class of problems where waves result from bistability. The interest for development, elaborated in discussion, is that waves propagate information faster than diffusion, and the proposed mechanism is generic and largely parameter independent.

Having captured the WNT response dynamics for the different cell lines, we can further model whether the differentiated cells become endoderm or mesoderm. To do so we assume that NODAL favours endoderm over mesoderm^4^, and we describe this as a branching probability from an intermediate mesendoderm state (Figure 7B), since we lack more quantitative data about the genetic network for mesendoderm specification. As a surrogate for NODAL signalling we used measured SMAD2 profiles at 24 hours for each cell line (Figure 5G and Figure M5 in Supplemental Methods). Figure 7E shows the comparison of model and data. The conversion of mesoderm to endoderm due to up regulation of NODAL is clearly visible in the comparison of CER1^-/-^ with wild type or E-CAD^-/-^.

## Discussion

Untangling the dynamics of early cell fate specification, and pattern formation during gastrulation remains a formidable challenge, especially during early human development. Using highly quantitative and standardized human gastruloids to model human gastrulation, we had previously shown the cascade of signaling begins with BMP4, which is sufficient to induce gastrulation and pattern all germ layers within 48 hours. We subsequently established that the mechanism underlying this BMP selforganization of gastrulation is based on two components: TGFβ receptors are only exposed to apical signals in a pre-pattern around the edge of the colony, and a Turing-like reaction-diffusion mechanism mediated by NOGGIN, which specifically in humans is induced directly by BMP4. We subsequently showed that the WNT pathway, operating downstream of BMP4, is necessary and sufficient for the mesoderm and endoderm layers and can self-organize its own pattern. In this work we have unraveled the molecular mechanism underlying this WNT-mediated self-organization.

This mechanism includes several elements. The first element is the “pre-pattern” of WNT sensitivity. Unlike the situation with BMP4 this pre-pattern is not imposed by polarity of receptor localization, but rather by β-CAT mechano-sensation via E-CAD, and the cytoskeleton, which is governed by tissue geometry. Given the importance of this pre-pattern for WNT patterning in a model human epiblast, how applicable might such pre-patterns be to gastrulation more generally? Gastrulation takes many forms across all animals, but a common theme is the correlation of WNT signalling with invagination^29^ (such as for example through the PS in amniotes or through a blastopore in lower orders). Invagination necessarily involves a breaking of geometric symmetry and a concentration of mechanical forces in an area of high curvature, and it is tempting to think that β-CAT mechanosensation is also involved here in either pre-patterning or reinforcing a WNT signal. Other notable examples of β-CAT mechanosensation as a critical element for WNT patterning include are found *Drosophila* gastrulation^30^, *Xenopus* embryonic explants^30^, and avian follicles^31^.

The second element in the WNT patterning process is the Turing-like activator-inhibitor pair of WNT and DKK1. In a manner similar to BMP4 which in species-specific manner induces NOGGIN, the WNT ligand also directly induces the expression of its own inhibitor: DKK1. In the mouse the anterior visceral endoderm (AVE) on the opposite side of the embryo from the PS is traditionally thought of as the major source of DKK1, with this inhibitor being produced in a WNT independent manner^32,33^. The expression of DKK1 by the AVE makes disentangling any intrinsic contribution of the epiblast to the position and control of the PS in the embryo difficult. Given that our synthetic epiblast achieves a stable pattern via the induction of DKK1 by WNT without external sources, however, we might expect to see DKK1 produced by cells in the PS. This is exactly what recent single cell RNA-seq data shows^34^. Similarly, results from studies in rabbit gastrulation show that DKK1 is also expressed in cells in the PS, especially in epiblast cells undergoing EMT^35^. While expression of activator and inhibitor on opposite sides of an embryo (such as in PS and AVE) seems logical and necessary, the Turing mechanism requires production of the inhibitor where the activator is highest, since the inhibitor must also spread rapidly to confine the activator.

We discover that the same is true for the CER1, the third major element in the WNT patterning process in our gastruloids. Despite also being thought of as an AVE product, single cell RNA-seq studies show that CER1 is also expressed in the mouse PS^34^. Separate rabbit studies confirm this^35^. In our human gastruloids we find that CER1 acts as a NODAL inhibitor and controls the balance between mesodermal and endodermal fates emerging from the back of the EMT wave. Thus E-CAD, DKK1, and CER1 are the primary determinants of WNT-induced PS patterning in artificial human epiblasts.

Our knockouts of CER1 and DKK1 in the gastruloids also reveal possible species specific differences between human, mouse, chick, and rabbit. For example, the mouse Dkk1 knockout does not broaden the PS as defined by Bra expression, though it does broaden the expression of a synthetic Wnt reporter^36^. This could suggest that PS formation in human is more dominated by WNT signals whereas in mouse the PS and Bra require other signals (such as NODAL) as well. Another difference between human and other studied species is that the knockout of CER1, or even the double knockout of CER1 and LEFTY1, does not lead to an expansion of the PS in our model system. This is surprising since in mouse it leads to multiple streaks^25^ and in the chick NODAL repression by Cerberus is thought to be the dominant repressor of ectopic streaks^37^. We hope that future comparative experiments will shed light on this apparent difference.

Our results for the DKK1 and CER1 knockouts have implications not just for the spatial control of the streak, but also for mesoderm versus endoderm cell fate decisions within the streak. We found a higher ratio of mesoderm to endoderm in DKK1^-/-^ cells, which is consistent with the reduction in the mouse of anterior endoderm in favour of mesoderm following the analogous knockout^36^. Conversely, removal of CER1 favors endoderm over mesoderm in our system, with upregulation of nuclear SMAD2 compared with wild type. Although less well known than the appearance of multiple streaks, the elimination of CER1 and LEFTY in the mouse embryo also results in a marked conversion of BRA cells to more NODAL regulated fates, such as anterior endoderm and axial mesoderm^25^. Based on these observations, as well as the result that WNT induces both DKK1 and NODAL in hESCs, we speculate that mammalian anterior endoderm is specified by a transient WNT3 pulse terminated by self-induced DKK1 and followed by up regulation of NODAL signalling. In fact, a transient application of WNT followed by its removal and the addition of ACTIVIN is precisely the most efficient and commonly used endoderm differentiation protocol used for hESCs^38,39^.

One of the more intriguing results to emerge from our investigation was the presence of a propagating EMT wave from the edge of our gastruloids towards the center. This wave is generated by downregulation of E-CAD by EMT and the negative regulation of WNT signalling by E-CAD, but also requires ACTIVIN/NODAL activity since SB treatment blocks the wave. Although it is harder to cleanly isolate in the embryo, there is evidence for cooperative EMT in maintaining and extending the PS, and arguments to this effect have been advanced in mouse and chick^40,41^. Traveling waves of activity have been seen in several other developmental contexts, for instance, calcium waves following fertilization or in large embryos that are presumably to synchronize tissues^42^. A wave of mitotic activity in frog extracts has also recently been observed and linked to bistability in the CDK system^43^. Why might a wave be useful for patterning? A wave is rationalized as way to spread information more rapidly than diffusion in a large system, and the hundred micron scale disk-shaped epiblast in rabbit and humans may require such a solution. Patterning via a wave may be a widespread mechanism for tissue patterning since it only requires a bistable system and some means for the favoured state to spread between cells.

Our gastruloids derived from hESC provide a platform where complex developmentally relevant patterning dynamics can be followed, quantified, and ultimately deconstructed. It thus provides a means to understand how embryonic development can be both so robust and yet flexible, and to ultimately apply this understanding to engineer systems to further regenerative medicine.

## Acknowledgements

hE-cadherin-pcDNA3 and hE-cadherin/Δβ-catenin-pcDNA3 were gifts from Barry Gumbiner (Addgene plasmids #45769 and #45772). This work was supported by grants R01 HD080699 and R01 GM101653, and NSF grant PHY 1502151 to E.D.S. We thank members of our lab for constructive comments, and we thank the staff of the Rockefeller Bio-Imaging Resource Center (BIRC) for their guidance in live imaging of the GLR reporter cell lines.

## Author Contributions

Conceptualization and writing, I.M., A.H.B., and E.D.S.; Experiments and construction of knockout lines performed by I.M.; All authors reviewed the manuscript.

## Author Information

Correspondence should go to E.D.S. or A.H.B. Material requests should be addressed to A.H.B.

## METHODS

### Cell Culture

All experiments used either the RUES2 hESC cell line, CRISPR/Cas9 edited cell lines based on this line, or CRISPR/Cas9 edited versions of the previously described RUES-GLR cell line^4^. hESCs were maintained in HUESM medium conditioned by mouse embryonic fibroblasts (MEF-CM) with additional 20 ng/mL bFGF. Mycoplasma testing was carried out before beginning experiments and again at regular 2-month intervals. Cells were grown on GelTrex (Invitrogen) coated tissue culture dishes (BD Biosciences, 1:40 dilution) for maintenance and expansion. Dishes were coated overnight at 4C and then incubated at 37C for at least 20 minutes before the cells were seeded onto the surface. Passaging was performed using Gentle Cell Dissociation Reagent (Stem Cell Technologies 07174).

### Micropatterned Cell Culture

Micropatterns were created using glass coverslips from CYTOO. The coverslips were first coated with 10 ug/ml laminin 521 (Biolamina) then diluted in PBS with calcium and magnesium (PBS++) for 3 h at 37°C. hESCs were dissociated with StemPro Accutase (Life Technologies) for 7 minutes and then washed once with growth media, washed again with PBS, and then resuspended in growth media with 10μM ROCK-inhibitor Y-27632 (Abcam) in 35mm tissue culture plastic dishes. For each coverslip 1×10^6^ cells in 2mL of media were used. After 1 h ROCK-inhibitor was removed and was replaced with standard growth media supplemented with Pen-Strep (Life Technologies). Cells were stimulated with the following ligands or small molecules 12 h after seeding: 100ng/mL WNT3A, 100ng/mL Activin-A, 10μM SB, 6μM CHIR.

### Immunocytochemistry

Cells were fixed in 4% paraformaldehyde for 20 minutes at room temperature, washed twice and stored in PBS. Permeabilization (30 minutes at room temperature), primary antibody incubation (overnight at 4C), and secondary antibody incubation (30 minutes at room temperature) were carried out with blocking buffer (3% donkey serum and 0.1% Triton X-100 in PB). After the primary incubation cells were washed 3 times with PBST (PBS + 0.1% Tween-20) for 30 minutes each. After secondary incubation cells were washed twice more with PBS, and then mounted on glass slides for imaging. All of the primary antibodies used are listed in Table 1.

**Table 1.** Antibody information

| Antigen | Antibody | Dilution |
| --- | --- | --- |
| Active β-CATENIN | Millipore 05-665 | 1:400 |
| BRACHYURY | R&D Systems AF-2085 | 1:300 |
| E-CADHERIN | Cell Signalling 3195 | 1:200 |
| FRZD7 | R&D Systems MAB1981 | 1:200 |
| LEF1 | Cell Signalling 2230 | 1:200 |
| LRP6 | R&D Systems 1505 | 1:100 |
| N-CADHERIN | BioLegend 350802 | 1:200 |
| SMAD2 | BD bioscience 610842 | 1:100 |
| SOX17 | R&D Systems AF-1924 | 1:200 |
| SOX2 | Cell Signalling 3579 | 1:200 |

### Microscopy and Image Analysis

All images used were acquired with a Zeiss Axio Observer using ZEN software and a 20×/0.8 numerical aperture (NA) lens. All image analysis was conducted using custom software written in Matlab. Images were first background subtracted and normalized and then stitched on a colony by colony basis. Scenes for background subtraction and normalization were acquired in the spaces between colonies where no cells were present. For segmentation of individual cells, we first used Ilastik classification to separate foreground from background. The classifier was trained for each experiment on the DAPI images of 4 randomly chosen colonies stitched colonies from that experiment. Once foreground and background were obtained, the DAPI channel was then filtered with a median and h-max filter and subtracted against a gradient of the image in order to identify the nuclei centers. These centers were then used as seeds for a watershed, against which the background mask was applied to obtain the final segmentation. Using this segmentation mask we then obtained average intensities for each cell of the nuclear markers in the other channels. For radial plots, the intensity of IF signal for each marker was normalized to the DAPI intensity, and these corrected singlecell expressions were then radially binned and averaged. The final radial profile represents the average of the indicated number of colonies. For clustering and classification, single-cell intensities were log transformed and then clusters were fit with a Gaussian mixture model (see Figure S4G for example).

### qPCR and RNA-seq

RNA was collected in Trizol at indicated time points from either micropatterned colonies or from small un-patterned colonies. Total RNA was purified using the RNeasy mini kit (Qiagen). qPCR was performed as described previously^3^, and primer sequences are listed in Table 2. RNA-seq is from a previously published data set^3^, and all raw data are available from the GEO database, accession number GSE77057.

**Table 2.**
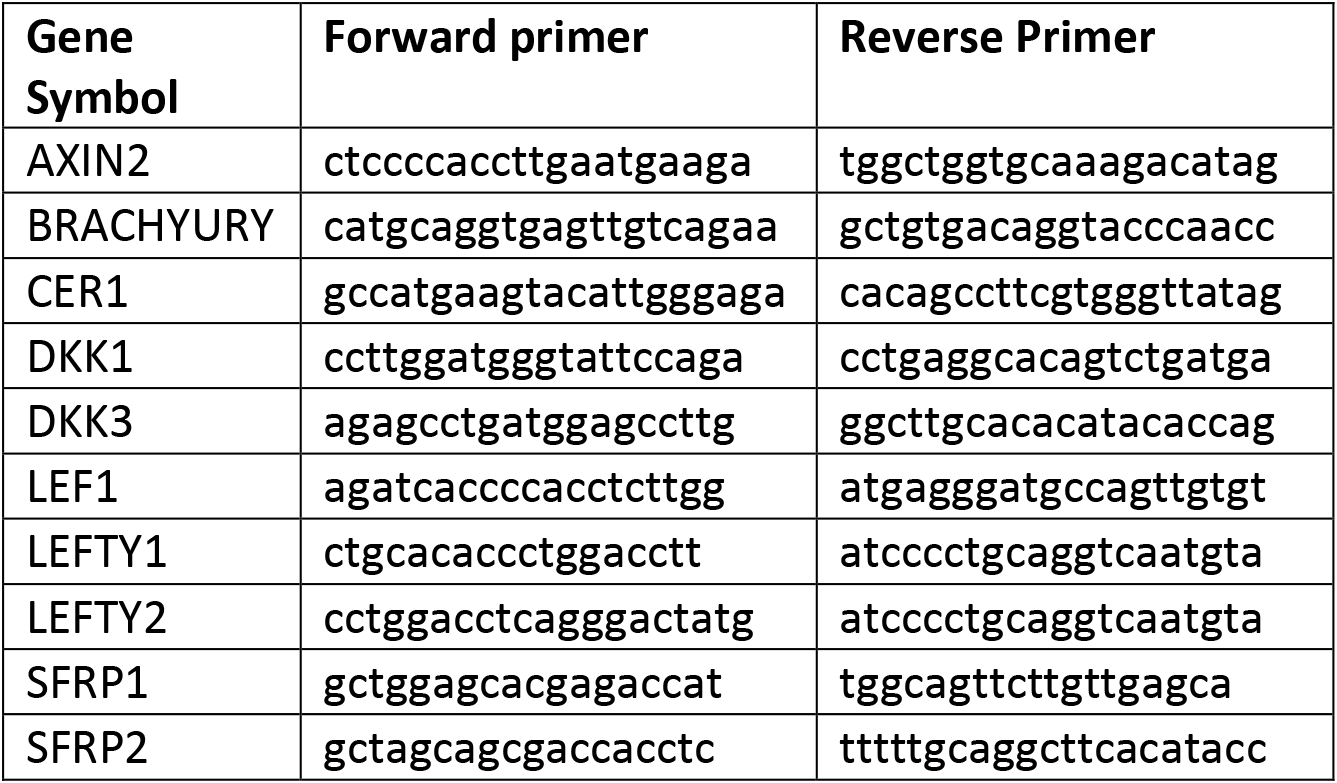
RT-QPCR Primer designs

### Transwell Experiments

We used Costar Transwell 24-well plates with 0.4μm pore-sized clear polycarbonate membrane inserts (Fisher Scientific 07-200-147). Membranes were coated with 10 μg/mL of laminin 521 (Biolamina) diluted in PBS++ for 3 hr, followed by washing 3 times with PBS++. Single cells were collected and seeded as per micropattern protocol. To image the membrane the transwell was removed from the multi-well plate after fixing and staining and placed on top of a coverslip.

### Generation of E-CAD insertion lines

Full length E-CAD and E-CAD lacking β-CAT binding domain were amplified via PCR from hE-cadherin-pcDNA3 and hE-cadherin/Δβ-catenin-pcDNA3 plasmids (Addgene plasmids #45769 and #45772) and inserted downstream of a pCAG promoter and puromycin resistance cassette flanked by 1kb homology arms for the AAVS1 safe harbour locus. This plasmid and TALENS targeting the AAVS1 site were then nucleofected into 1×10^6^ pluripotent E-CAD^-/-^ cells using the B-016 setting on an Amaxa Nucleofector II (Lonza). Nucleofected cells were then plated as per maintenance conditions, but supplemented with 10uM ROCK-inhibitor. Selection for both puromycin commenced after 2 days, and ROCK-inhibitor was maintained until colonies reached adequate size (typically 8-16 cells per colony). To derive pure clones, individual colonies were picked in an IVF hood with a 20uL pipette tip and seeded into separate wells with growth media and ROCK-inhibitor. Once successfully established, each clone was assayed functionally for brightness and homogeneity of the overexpressed E-CAD.

### Generation of knockout lines

The CRISPR/Cas9 system was used to generate all knockout lines. Three rules of reproducibility and quality control were applied: (i) at least two independent clones for each gene clonal lines were isolated and studied in parallel; (ii) lack of off-target effects was assessed by qPCR; and (ii) their ability to maintain the pluripotent state as assessed by expression of NANOG, OCT4, and SOX2. For the E-CAD, DKK1, and CER1 knockouts one clone of each was also assessed for chromosomal integrity by karyotyping. The sgRNA target for each gene is listed in Table 3. A list of all knockout lines is given in Table 4. The sgRNAs were cloned into a pX330 plasmid^44^ that we modified to co-express a puromycin-2A-EGFP cassette, as this strategy gave us a higher percentage of successfully targeted clones. Transfection was carried out using the B-016 setting of a Nucleofector II instrument and using the Cell Line Nucelofector Kit L (Lonza). Transfected cells were immediately seeded in ROCK-inhibitor on GelTrex coated culture dishes, and puromyacin was added after 24 h for 24 h. Cells were then passaged as single cells using Accutase (Stem Cell Technologies) and sparsely seeded to facilitate picking individual clones. Clones were handpicked with a 20uL pipette tip and once expanded genomic DNA from each clone was extracted with the DNeasy Blood & Tissue kit (Qiagen). The locus for each targeted gene was then PCR-amplified using the primers listed in Table 4, and submitted for Sanger sequencing. The resulting chromatograms for each clone were decomposed using the TIDE webtool^45^ http://tide.nki.nl. Only clones that showed a high probability for both alleles of the gene of interest having a missense mutation leading to a premature stop codon were kept. The chromatograms and TIDE decomposition for these clones are shown in Figure S6.

**Table 3.**
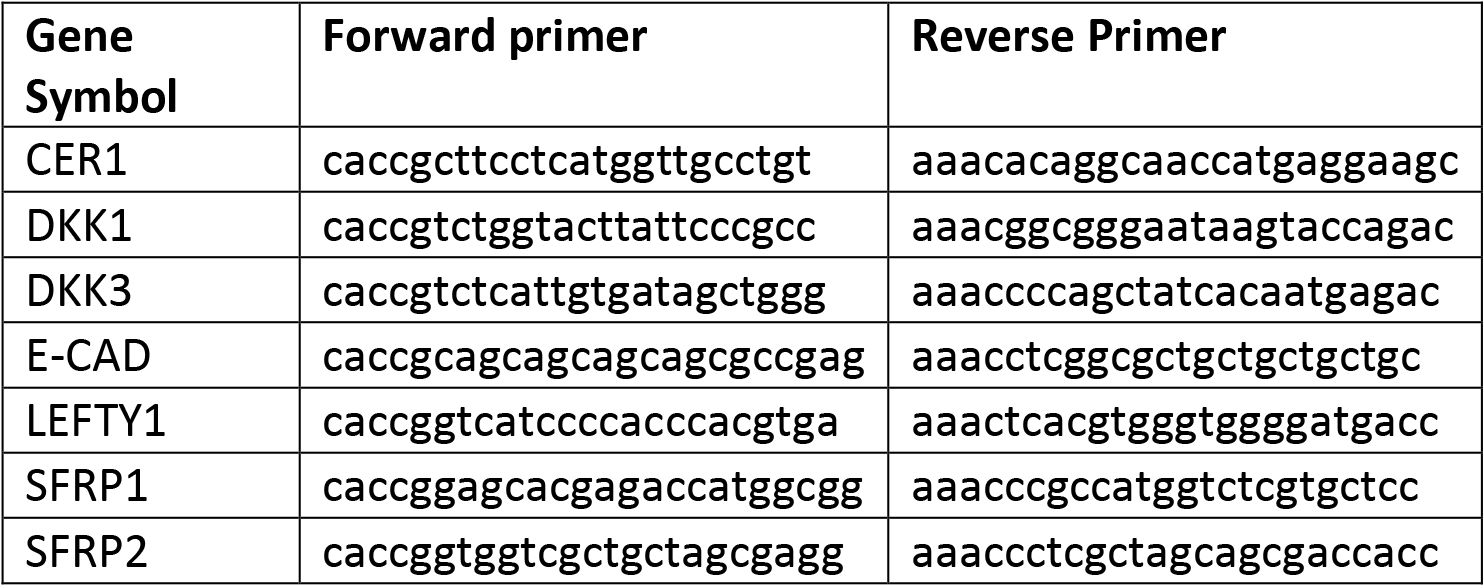
sgRNA designs

**Table 4.** Knockout lines

| Target | Separate clones |
| --- | --- |
| RUES2:DKK1<sup>-/-</sup> | 3 |
| RUES2-GLR:DKK1<sup>-/-</sup> | 2 |
| RUES2:DKK3<sup>-/-</sup> | 2 |
| RUES2:E-CAD<sup>-/-</sup> | 2 |
| RUES2:CER1<sup>-/-</sup> | 1 |
| RUES2:LEFTY1<sup>-/-</sup> | 1 |
| RUES2:SFRP1<sup>-/-</sup> | 1 |
| RUES2:SFRP2<sup>-/-</sup> | 1 |
| RUES2:SFRP1<sup>-/-</sup>SFRP2<sup>-/-</sup> | 1 |
| RUES2:LEFTY1<sup>-/-</sup>CER1<sup>-/-</sup> | 1 |

---

**Supplemental Figure 1.**
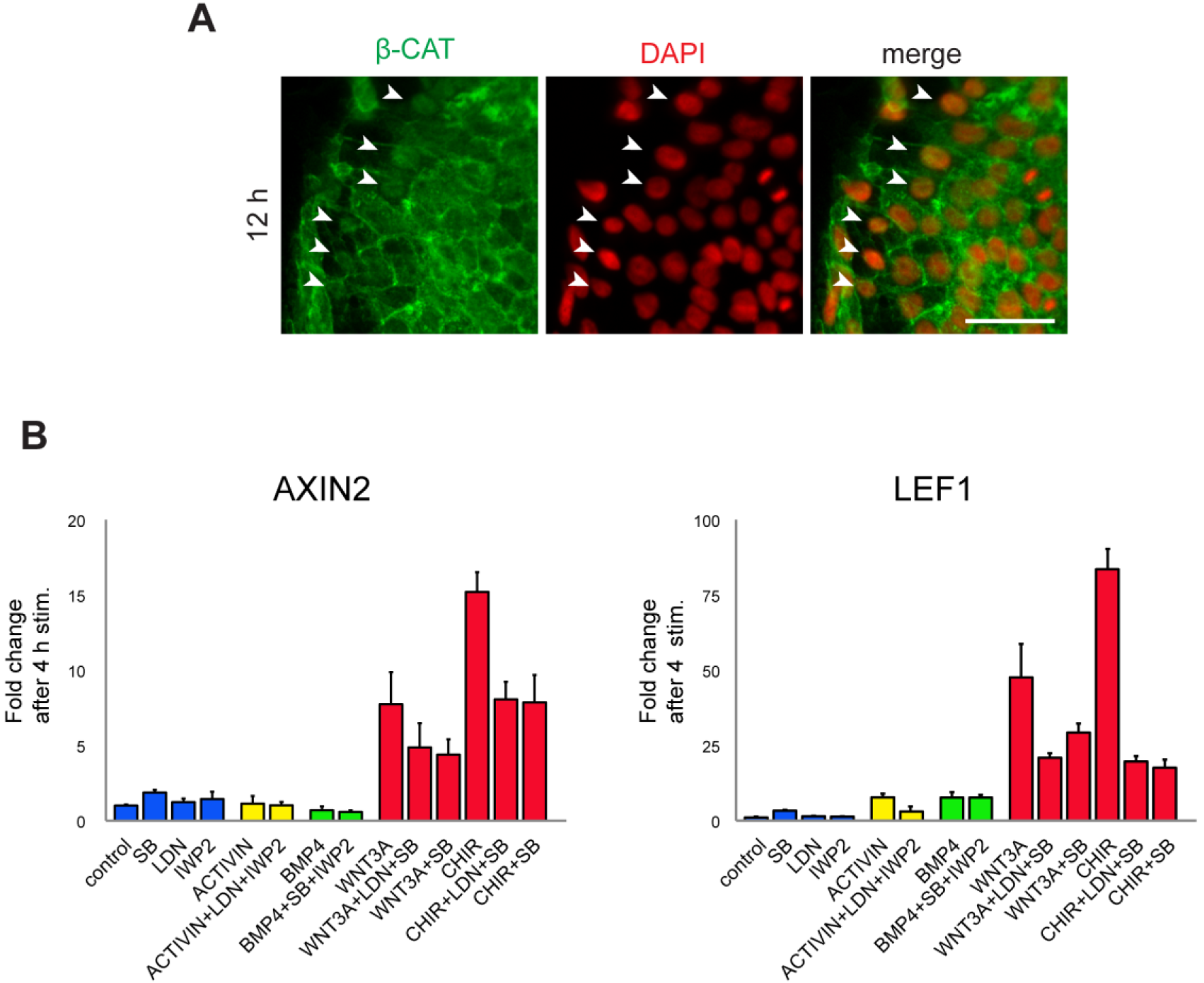
WNT3A response markers. (A) Inset of an abnormally low density micropattern colony stimulated with WNT3A and fixed and stained at 12h for active β-CATENIN. Even at this low density (seeded with 6×10^5^ cells, stimulated 2 hours after seeding) an increase in β-CAT in the nucleus is only visible in loosely connected cells on the colony edge (arrows). Nuclear β-CATENIN in cells away from the periphery or at higher density (including standard low density conditions) is not easily observed, largely due to signal from membrane-bound or cytoplasmic regions. Scale bar, 50μm. (B) qPCR of AXIN2 and LEF1 as a function of inputs arrayed on x-axis. The results show that the LEF1 and AXIN2 response is dominated by WNT3A, though synergism with the NODAL pathway is significant as well, as can be seen by comparing WNT3A or CHIR with WNT3A+SB or CHIR+SB. Thus LEF1 and AXIN2 can be used as proxies for early WNT3A response. Also note that LEF1 gives a greater positive signal than AXIN2, reaching 84-fold induction compared with 15 fold induction for CHIR condition.

**Supplemental Figure 2.**
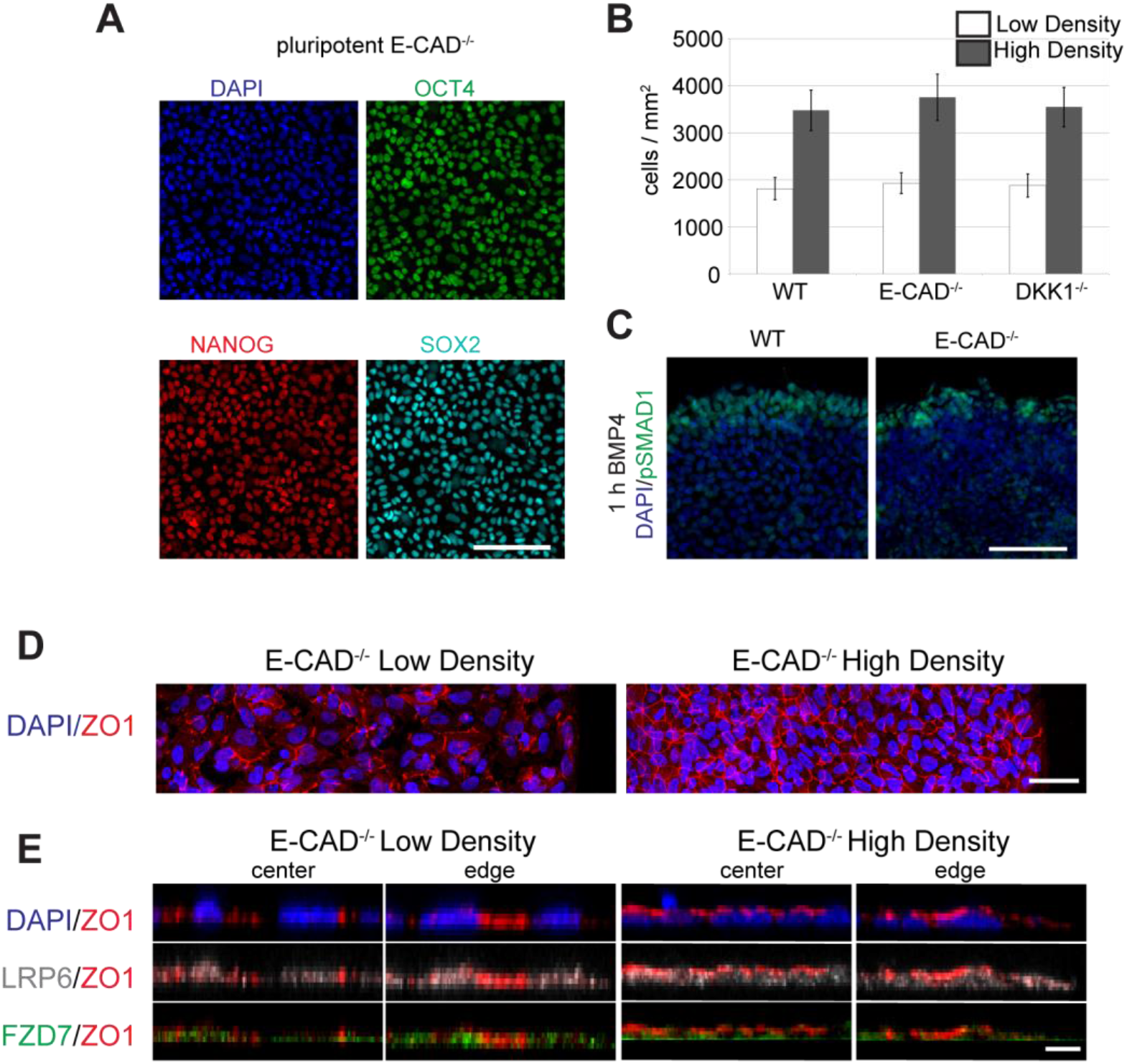
Epithelial integrity is preserved in E-CAD^-/-^ cells. (A) E-CAD^-/-^ cells maintain pluripotency markers even with continual passaging (i.e. >20 passages). Scale bar, 100μm. (B) E-CAD^-/-^ cell seeding efficiency and growth is similar to wild type and DKK1-/- cell lines. (C) Test of epithelial integrity and BMP4 response. Edge of high density wild type and E-CAD^-/-^ micropatterns stimulated with BMP4 and fixed and stained for pSmad1 after 1h. As in the wild type, pSMAD1 expression is restricted to the periphery in E-CAD^-/-^ colonies. Scale bar, 100μm. (D) Maximum intensity projection of ZO1 and DAPI in low density and high density E-CAD^-/-^ micropatterns immediately prior to stimulation. The network of tight junctions is the same as in the wild type (Figure 1C). Scale bar, 50μm. (E) Cross-sections showing the apical-basal position of WNT receptors relative to DAPI and ZO1. Result is the same as for wild type micropaterns (Figure 1D). Scale bar, 10μm.

**Supplemental Figure 3.**
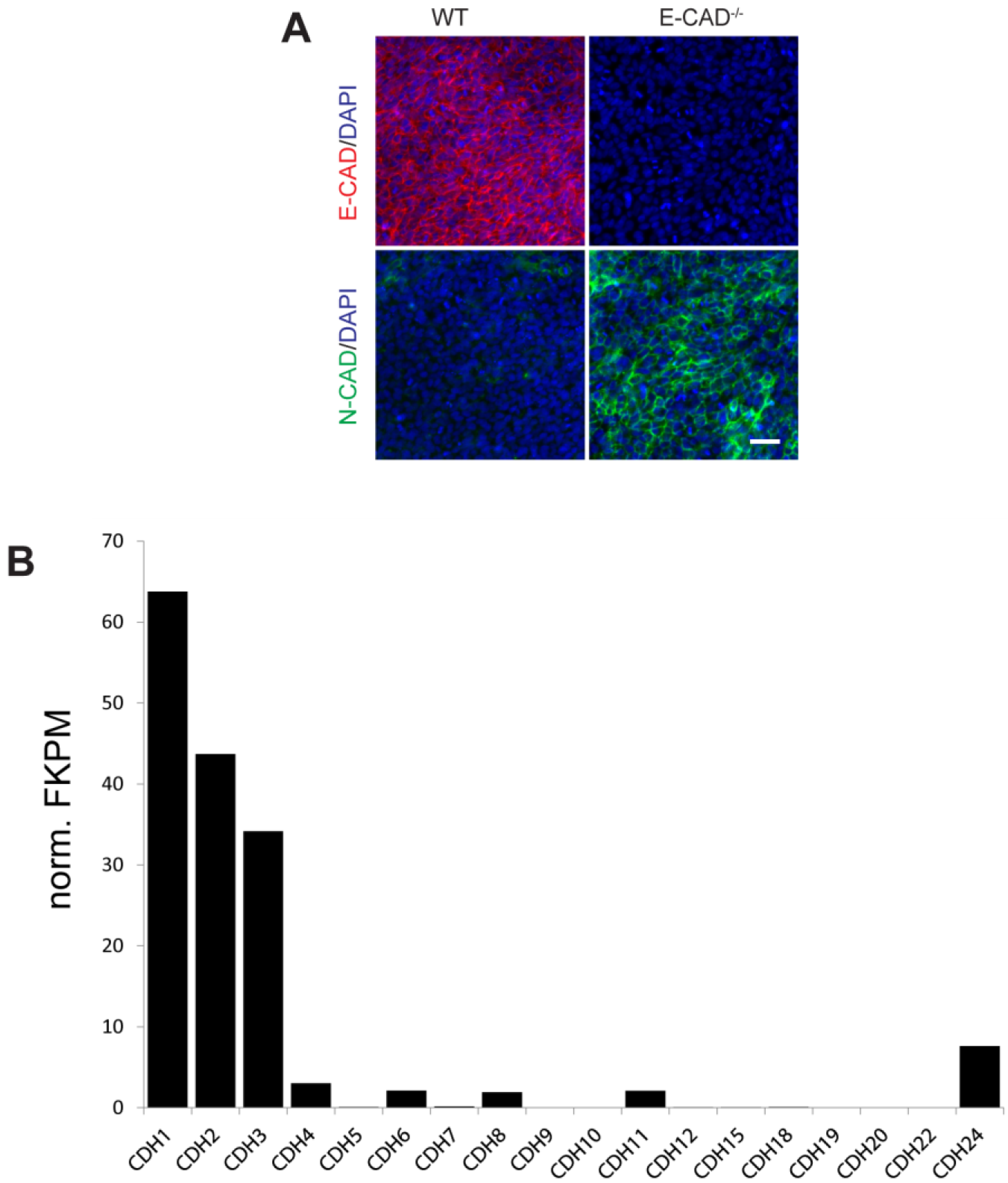
A change in the protein expression profile of N-CADHERIN occurs in the E-CAD^-/-^ cell line. (A) RNA-seq profiling for all classic cadherins in pluripotent hESCs. Since the E-CAD (CHD1) knockout is lethal in mouse at the post-compaction stage, we were somewhat surprised at the lack of an E-CAD^-/-^ pluripotent phenotype (Supplemental Figure 2). However both N-CADHERIN (N-CAD) and P-CADHERIN (CHD2,3) are substantially expressed at the mRNA level in hESCs in pluripotency. (B) Stain for N-CAD and E-CAD in wild type and E-CAD^-/-^ in unpatterned pluripotent colonies. Antibody stain for E-CAD confirms that gene is knocked out in E-CAD^-/-^ cells. More interestingly, while N-CAD is barely visible in wild-type cells, N-CAD is highly expressed and membrane localized in E-CAD^-/-^ cells. Thus there is a reservoir of N-CAD message in hESC that is only expressed in the absence of E-CAD, which may be a consequence of the same pathway that up regulates N-CAD during EMT when the transcription of E-CAD is abrogated by SNAIL. Interestingly, other research has shown that the artificial replacement E-CAD protein by N-CAD protein in the mouse intestine after gastrulation showed that N-CAD could fulfill the structural role of E-CAD, but the replacement lead to an up-regulation of WNT signalling that is also consistent with our findings^46^. Scale bar, 50μm.

**Supplemental Figure 4.**
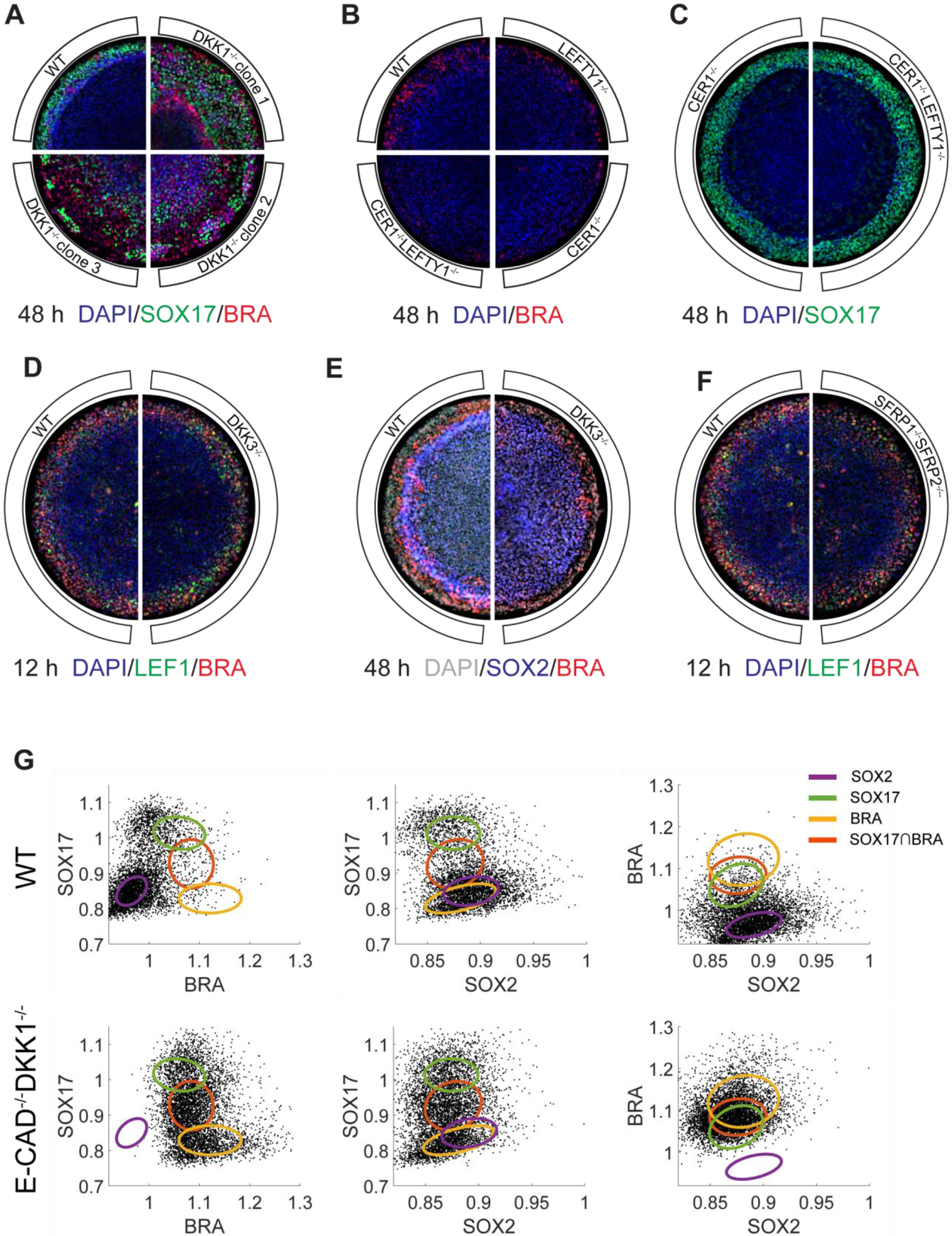
Other inhibitor CRISPR knockout line and clones. (A) Other DKK1^-/-^ clonal lines with different frameshift mutations also exhibit the same micropattern phenotype when stimulated with WNT3A at high density and fixed and stained after 48 hours. (B) No discernible difference in SOX17 expression at 48h between CER1^-/-^ and CER1^-/-^LEFTY1^-/-^ micropatterns under WNT3A stimulation. (C) LEFTY1^-/-^ micropatterns show no discernible difference with wild type micropatterns in number of BRA cells. However, both CER1^-/-^ and CER1^-/-^LEFTY1^-/-^ (with a different CER1 frameshift mutation) show similar phenotype in having fewer BRA cells. Thus CER1^-/-^ and not LEFTY1^-/-^ is the main NODAL inhibitor during WNT induced patterning. (D) and (E) No discernible difference at 12h or 48h between wild type and DKK3^-/-^ micropatterns under WNT3A stimulation. (F) No discernible difference at 12h between wild type and SFRP1^-/-^SFRP2^-/-^ micropatterns under WNT3A stimulation. (G) Example scatterplots of wildtype and E-CAD^-/-^DKK1^-/-^ micropatterns data showing clustering into the Gaussian mixture model used for Figure 5E. Here the 3D Gaussians are projected into 2D for better visualization.

**Supplemental Figure 5.**
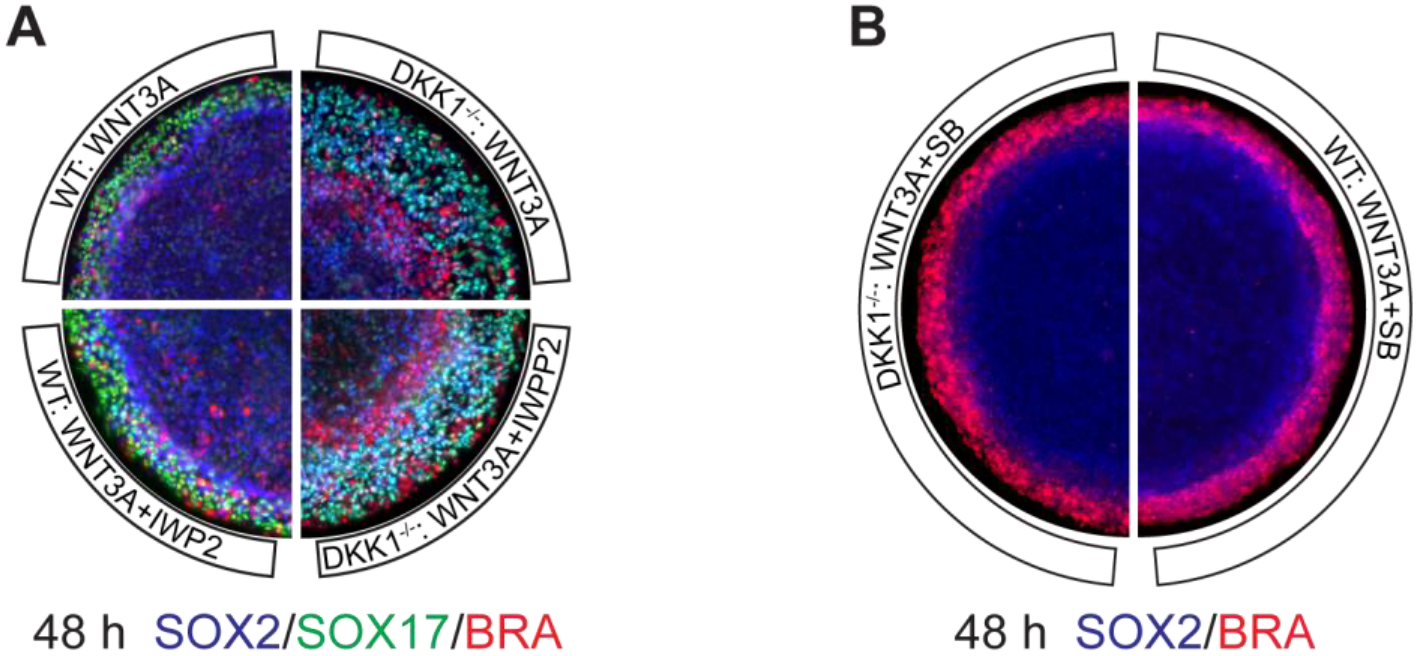
Endogenous WNT signalling has little effect on patterning. (A) Wild type and DKK1^-/-^ high density micropatterns stimulated with either WNT3A or WNT3A+IWP2 for 48h and stained for BRA, SOX2, and SOX17. No significant differences between wild type IWP2 and non-IWP2 or DKK1^-/-^ IWP2 and non-IWP2 stimulated colonies were observed. (B) Wild type and DKK1^-/-^ high density micropatterns stimulated with WNT3A+SB for 48h and stained for BRA and SOX2.

**Supplemental Figure 6.**
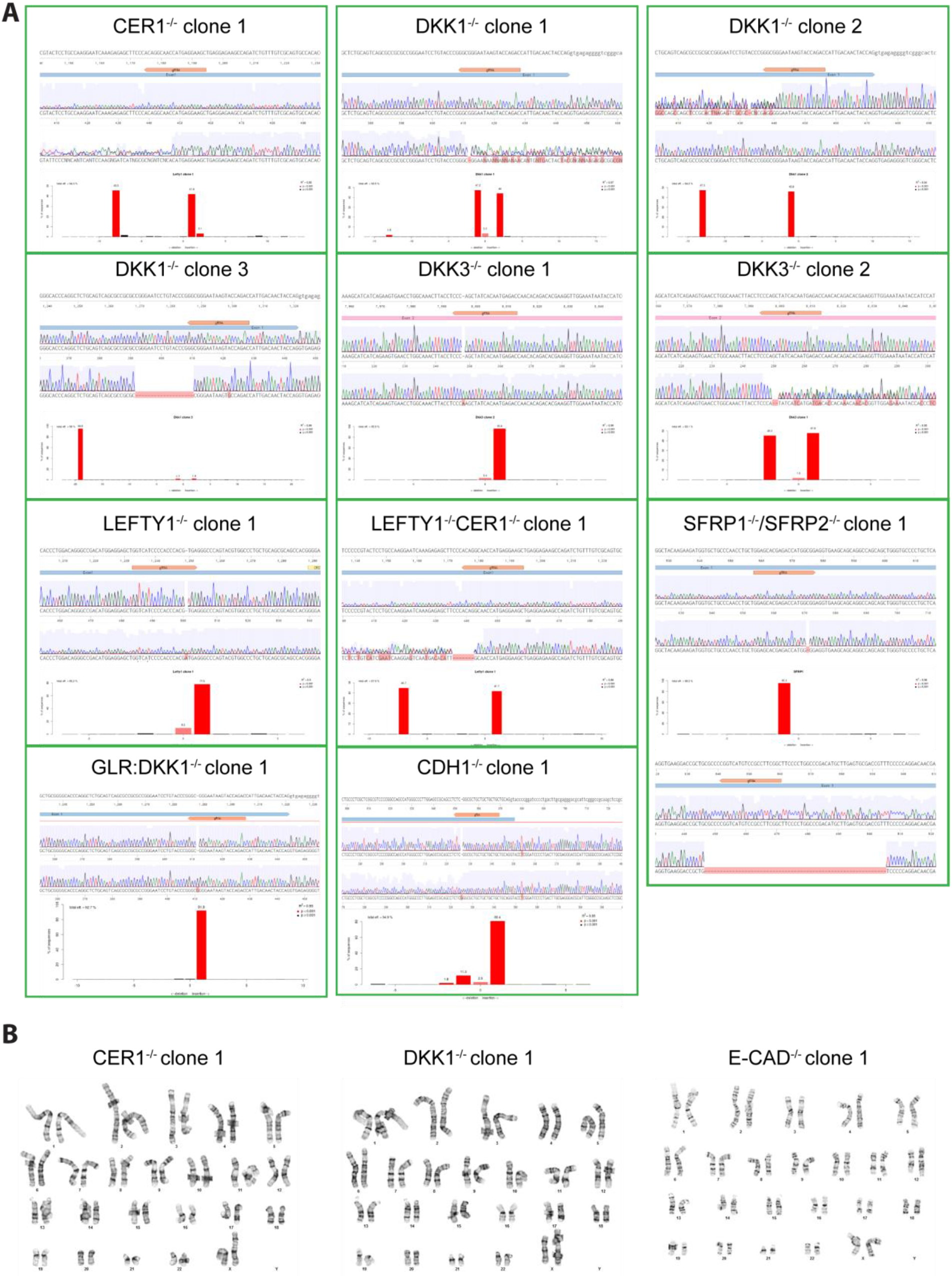
Confirmation of CRISPR knockouts. (A) Sequencing to confirm knockout of WNT and NODAL inhibitors. The resulting chromatograms for each clone is shown, as well as the decomposition using the TIDE webtool^45^ http://tide.nki.nl. Only clones that showed a high probability for both alleles of the gene of interest having a missense mutation leading to a premature stop codon were considered validated. Note that due to the low rate of successful targeting of the LEFTY1 locus, the LEFTY1^-/-^CER1^-/-^ clone 1 cell line was built by additionally knocking out CER1 from the LEFTY1^-/-^ clone 1 cell line. Also, the TIDE decomposition for SFRP2 is not shown as the tool cannot handle indels as large as 59bp, which is the amount that can be seen to be deleted in both alleles in the chromatogram. (B) The E-CAD^-/-^ clone 1, DKK1^-/-^ clone 1, and CER1^-/-^ clone 1 cell lines were karyotypically normal.

